# Ectopic overproduction of cell wall glucan through membrane perturbation by an antifungal peptide theonellamide A in fission yeast

**DOI:** 10.64898/2026.04.23.720496

**Authors:** Kensuke Nakao, Vanessa S. D. Carvalho, Akifumi Suganaga, Masako Osumi, Masato Tokukura, Hideaki Kakeya, Akihisa Matsuyama, Yoko Yashiroda, Shigeki Matsunaga, Juan C. G. Cortés, Minoru Yoshida, Juan C. Ribas, Shinichi Nishimura

## Abstract

Ergosterol has multiple functions in filamentous fungi and yeasts, although only a part of the functions seems to be understood. An antifungal peptide, theonellamide A (TNM-A) induces drastic morphological changes in fission yeast cells by targeting plasma membrane ergosterol. TNM-A induces overproduction and ectopic accumulation of cell wall glucan at both growing tips and septum through a yet unknown mechanism. Here we show that TNM-A treatment causes accumulation of 1,3-β-glucan at cell-polarity sites, not by increased activity of 1,3-β-glucan synthase, but by an increased, persistent localization of the glucan synthase enzymes. Screening based on subcellular localization of proteins at periphery or polarity sites suggested the involvement of the Rho family GTPase Cdc42. In agreement, TNM-A induced both activation of Cdc42 and enhancement of membrane trafficking of glucan synthase enzymes. In conclusion, our chemical genetics analyses using TNM-A suggest that membrane ergosterol regulates the activity of Cdc42, which further regulates the localization of glucan synthases and cell wall biosynthesis.

**Highlights (four sentences):**

- Thenoellamide A (TNM-A) induces an ectopic overproduction of cell wall glucan.
- TNM-A treatment causes increased, persistent localization of glucan synthases at the cell tips and septum.
- TNM-A activates Cdc42 and upregulates membrane trafficking of glucan synthases.
- Ergosterol is involved in proper activation/inactivation of Cdc42.

## INTRODUCTION

Sterols are the essential components in the eukaryotic cell membranes [1, 2]. Its concentration varies from organelle to organelle, while it accounts for 20-50 mol% in the plasma membrane. Sterols regulate the physical properties of the membrane such as fluidity to form membrane microdomains, while they have been reported to be important for structures and functions of membrane proteins [3]. However, we have not fully understood the function of sterols partly due to the lack of a standard experimental approach. Genetic manipulation of the lipid-biosynthesis genes often induces unexpected, secondary metabolic changes, while chemical modification of lipids to yield chemical probes often results in the loss of the physical properties of the parent lipid molecules. In contrast, chemical genetics utilizing bioactive compounds that exhibit unique phenomena by binding to specific lipid molecules is sometimes effective for revealing hidden lipid functions [4].

Theonellamides (TNMs) are marine-sponge derived bioactive peptides, which bind to 3β-sterols including cholesterol and ergosterol. They bind to sterols at the shallow area of the lipid membranes [5], favor sterols in the liquid disordered domains and change the fluidity [6], and modulate the membrane curvature and vesicle sizes [7]. In contrast to other sterol-targeting molecules such as amphotericin B and filipin, TNMs induce neither acute toxicity nor cell lysis [8, 9]. In the fission yeast *Schizosaccharomyces pombe*, TNMs induce an overproduction of cell wall material, which is dependent on Bgs1 and Rho1 GTPase, suggesting that TNMs induce overproduction of 1,3-β-glucan [10]. Bgs1 is one of the four essential catalytic subunits of the enzyme 1,3-β-glucan synthase (GS) in fission yeast, and Rho1 was shown to be the regulatory subunit required for the activity of 1,3-β-glucan synthases [11, 12]. However, the composition of the overproduced cell wall material and the molecular mechanism underlying the overproduction have not been clarified.

Cdc42 is a small Rho GTPase that is conserved among eukaryotes and regulates cell polarity [13]. In fission yeast, Cdc42 regulates actin skeleton formation and membrane trafficking to establish the cell polarity [14]. For example, Cdc42 regulates polarized transport of Bgs1, which suggests that it also regulates cell wall biosynthesis [15]. Cdc42 activity is regulated by guanine nucleotide exchange factors and GTPase-activating proteins [16]. The regulatory events occur at the inner leaflet of the plasma membrane[17], but how membrane lipids are involved in that is not yet understood.

In this study, we show that TNM-A induces an increased, persistent localization of both β-glucan synthases (βGSs) and α-glucan synthase (αGS) Ags1/Mok1, and thus a cell wall glucan overproduction at cell tips and septum. Proteome analyses based on protein subcellular localization in cells treated with TNM-A unveiled the accumulation of proteins that function in the membrane trafficking, at sites of active cell wall biosynthesis. In fact, TNM-A activated Cdc42, by which GS seems to be accumulated at cell-polarity sites. Our results suggest that membrane ergosterol is involved in the timely activation of Cdc42, thus controlling proper cell wall biogenesis.

## RESULTS

### Accumulation of ectopic cell wall material at cell polarity sites (Figure 1, Figure S1-3)

During cytokinesis fission yeast generates a septum, by which a mother cell divides into two daughter cells. The septum is visualized by a fluorescent dye, calcofluor white (Cfw), which preferentially stains linear 1,3-β-glucan, one of the glucans in this organism. We previously reported that TNM-F, a related congener of TNM-A which is used in this study (Figure 1A), induces overproduction of cell wall at cell tips and septum which was visualized by Cfw fluorescence [10]. In this study we first analyzed both cell growth and morphology in the presence of increasing concentrations of TNM-A. Cell growth inhibition was observed at higher concentrations of TNM-A (1-10 μg/ml), while the lower concentration of TNM-A (0.5 μg/ml) exhibited partial inhibition (Figure S1A). The thick cell wall phenotype was observed under fluorescence microscopy at these concentrations (Figure S1B). The phenotype was attenuated at 5-10 μg/ml, when compared with concentrations of 1-2 μg/ml, likely due to the membrane damage that is observed at higher concentrations of TNM-F [10]. In this study, we applied a concentration of 2 μg/ml to examine the effect of TNM-A in cell morphology.

**Figure 1.**
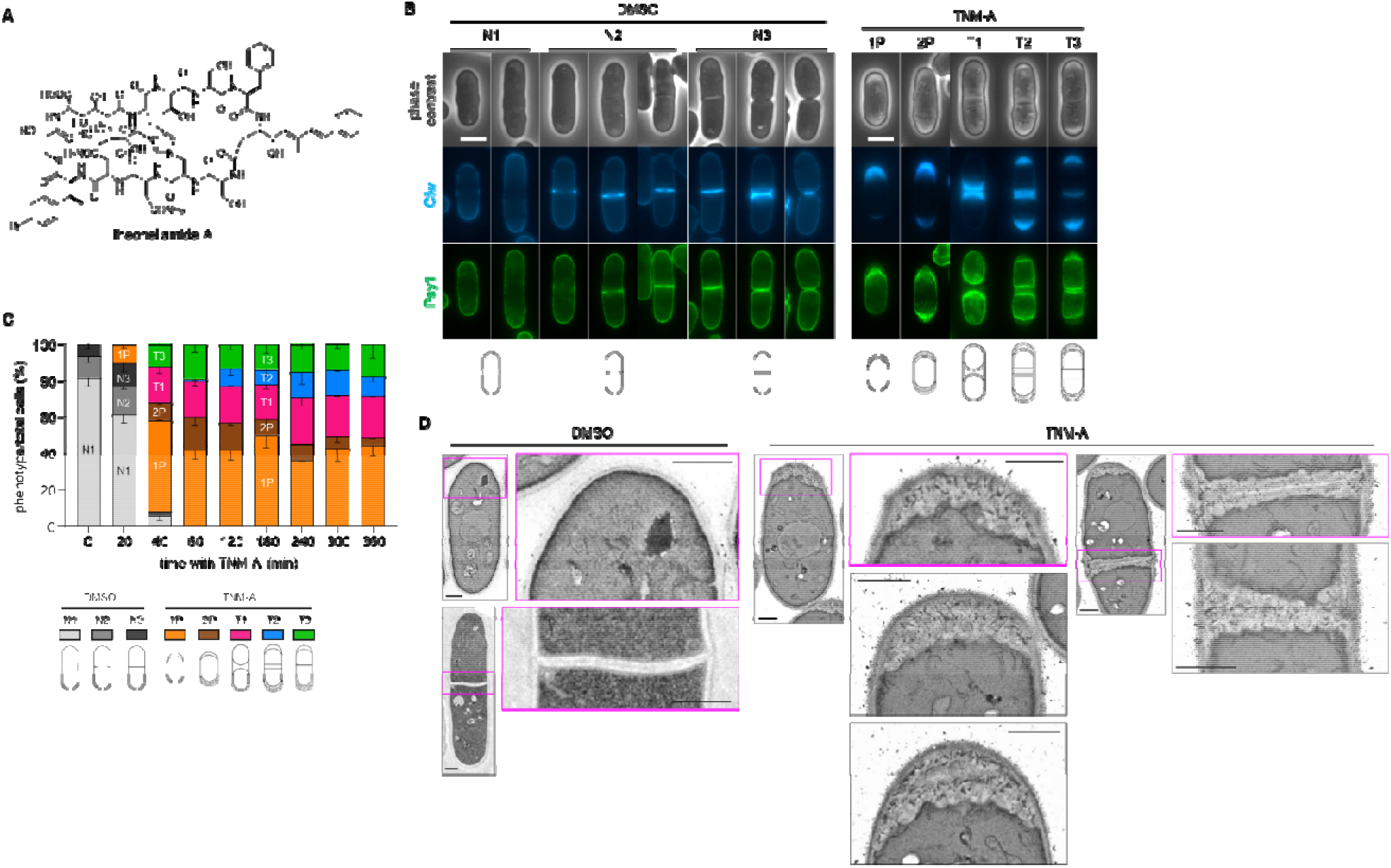
Thick ectopic cell wall at cell tips and septa generated by TNM-A. (A) Chemical structure of TNM-A. (B) Accumulation of 1,3-β-glucan by TNM-A, observed under fluorescence microscopy. Cells expressing GFP-Psy1 (a plasma membrane marker) were cultivated in YES at 28 °C, without or with 2.0 µg/mL TNM-A for 4 hours. Images of phase contrast, and Cfw and GFP fluorescence microscopy, are shown. Schematic representation of the thick ectopic cell wall phenotype for each cell phenotype with TNM-A during either interphase (1P, and 2P) or cytokinesis (T1, T2, and T3) is shown. Scale bar, 5 μm. (C) Time course analyses of the cellular morphologies over 6 hours with TNM-A. Schematic representation of the three types of normal cells with DMSO (N1, N2, and N3), and of the thick ectopic cell wall phenotype for each cell phenotype with TNM-A as in B (1P, 2P, T1, T2, and T3), is shown. Data represent the mean ± SD (4,129 cells in total). (D) Cell wall ultrastructure at the cell tip and septum of untreated cells (DMSO) or altered accumulation of ectopic cell wall material growing inwards either at the cell tip or septum of treated cells (TNM-A). Cells were treated with DMSO or TNM-A for two hours and processed for transmission electron microscopy (TEM). Areas with boxes are enlarged in the right panels. Scale bar, 1 μm.

In contrast to the control cells (Figure 1B, DMSO), after 4 hours of TNM-A treatment all the cells showed a phenotype of thick cell wall either at the cell tips, septum, or both. Cells could be classified into several groups. First, depending on the Cfw fluorescence in dividing cells, there are three types of cells with different septum thickness, named T1, T2, and T3 cells. T1 cells exhibited thick septa and no intense Cfw signal at cell tips, while T2 cells had thick septa and intense Cfw signal at cell tips. T3 cells showed a normal septum and intense Cfw signal at cell tips (Figure 1B, TNM-A). The septum thickness in T1 and T2 cells grew in a time-dependent manner, suggesting the continuous biosynthesis of glucan in these abnormally thick septa (Figure 1C and Figure S2). Second, depending on the Cfw fluorescence in the cell tips of non-dividing cells, two more cell types (1P and 2P) were observed: 1P cells displayed an increased Cfw fluorescence in one of the cell tips, while 2P cells were strongly stained by Cfw in both cell tips (Figure 1B). TNM-A induced glucan synthesis in unusual amounts at both septum and cell tips. Cells with normal morphology disappeared after 1 h treatment with TNM-A (Figure 1C).

To observe the ultrastructure of the thick cell wall, we analysed the cell morphology under both scanning electron microscopy (SEM) and transmission electron microscopy (TEM). In the SEM observation, the cell wall surface structure was indistinguishable between control cells and compound-treated cells (Figure S3). In the TEM analyses, drastic changes by TNM-A were observed. The cell wall forms a thin, uniform layer under DMSO-treated conditions (Figure 1D). In TNM-A-treated cells, ectopic materials with low electron density, which correspond to the network of glucans, were observed at the septa and cell tips, while the cytoplasmic surface is bumpy. Electron-dense materials, perhaps corresponding to glycoproteins, are scattered in disorder within the accumulated glucan. Since there were no notable changes in the cell surface structure and cell length, it was thought that abnormal ectopic cell wall synthesis occurred, while cell wall weakening and remodelling to permit cell wall expansion is blocked.

### Increased biosynthesis of glucan by TNM-A (Figure 2, Figure S4-6)

The cell wall of *S. pombe* consists of three types of polysaccharides, α-glucan, β-glucans, and galactomannan. We investigated the fission yeast cell wall with TNM-A biochemically. First the uptake of ^14^C-glucose into the cell wall was measured and compared to the cell (Figure 2A, Figure S4). ^14^C-glucose was taken up in a time-dependent manner in the presence of TNM-A, indicating that the synthesis of polysaccharides is increased and accumulated by the compound. Next the composition of the cell wall was investigated (Figure 2B, Figure S4). Among three types of polysaccharides, the percentage of β-glucan, specifically that of 1,3-β-glucan, but not that of 1,6-β-glucan, increased in a time-dependent manner. The cellular content of 1,3-β-glucan drastically increased from 18% (control cells) to 36 and 39% (4 h and 6 h treatment of TNM-A, respectively). These results are consistent with the fluorescence microscopy analyses using Cfw, which stains mainly linear 1,3-β-glucan. The second major component, α-glucan, also increased in the presence of TNM-A: 10% (control cells) to 15% (4 h treatment of TNM-A), while it decreased to 13% after 6h with TNM-A. The amount of the other cell-wall component, galactomannan, did not show increase both in the biochemical and cytological analyses (Figure 2B, Figures S4, and S5). These biochemical data revealed that the composition of the thick cell wall generated by TNM-A treatment is different from that of the normal cell wall: the increase in the amount of 1,3-β-glucan is particularly drastic.

**Figure 2.**
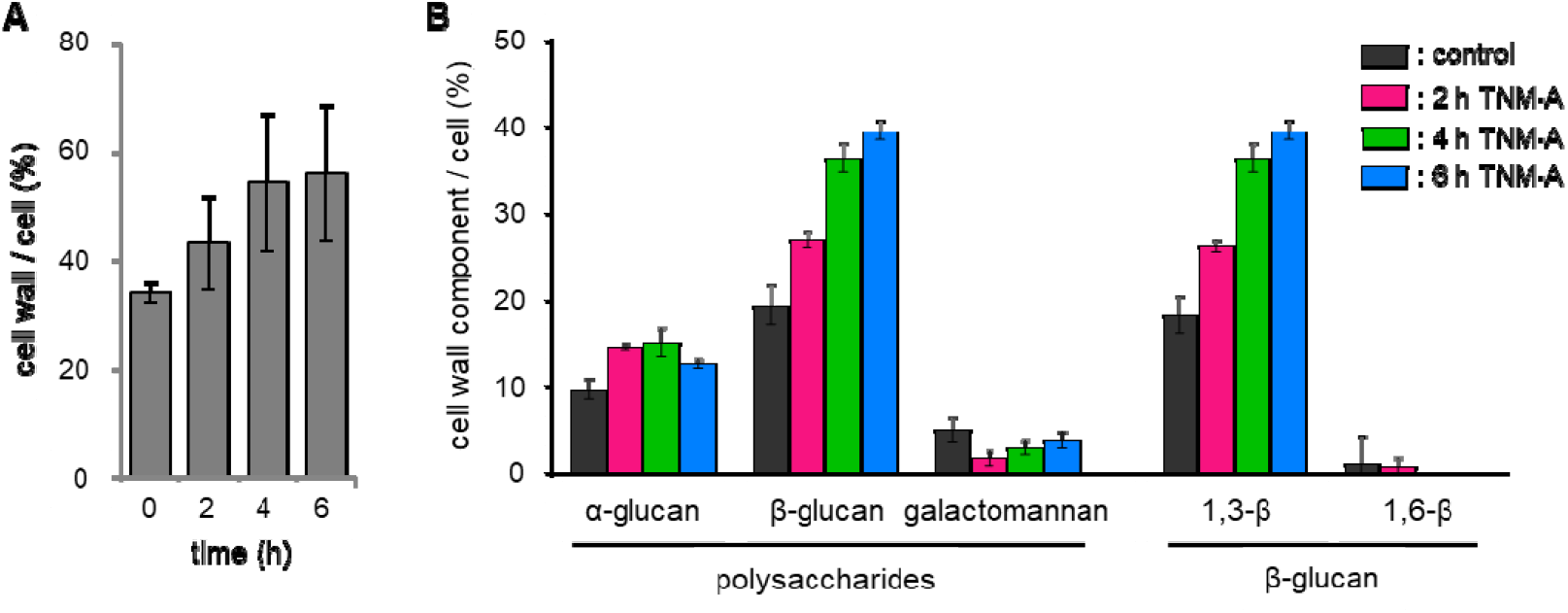
Increased biosynthesis of cell wall glucan by TNM-A. Cell wall amount (A) and cell wall composition (B) of *S. pombe* cells treated with TNM-A for 2, 4 and 6 hours are shown. Cells were incubated in YES medium at 28 ºC under shaking with TNM-A at 2.0 µg/mL and ^14^C-glucose for 6 hours prior addition of TNM-A. The cell walls were isolated to quantify their radioactivity with respect to the cell (A). The isolated walls were subjected to different treatments to fractionate and quantify their components (B). Data represent the mean ± SD (n = 4 independent experiments).

There are two possible explanations for the increased amount of 1,3-β-glucan in TNM-A-treated cells: the GS activities were upregulated or the amount of the GS proteins was increased at sites of active growth. The first possibility was examined by testing the GS activity in two ways (Figure S6). In the first experiment, cells were treated with TNM-A *in vivo* during increasing periods of time, and then the membrane fractions were prepared to measure the GS activity. The *in vivo* treatment of TNM-A did not increase the GS activity, instead, treatment of cells with the compound gradually decreased the GS activity to less than 50% activity after 4 h treatment (Figure S6A). In the second experiment, the GS activity of the membrane fraction was measured in the *in vitro* presence of increasing concentrations of TNM-A in the reaction. The compound reduced the GS activity, although its effect was moderate, with a maximum inhibition of 25% that did not further increase with higher concentrations of the compound (Figure S6B). These GS activity tests indicate that TNM-A does not directly activate the enzymatic activity but likely increases the amount and timing of GS proteins at cell tips and septa.

### Increased, persistent localization of Bgs and Ags1 proteins at cell polarity sites (Figure 3, Figure S7-11)

The localization of GSs was analysed under fluorescence microscopy. Fission yeast has three characterized glucan synthases, Bgs1, Bgs4, and Ags1. All three proteins are known to be localized at growing areas: the growing tips during interphase and site of septation during cytokinesis [12, 18, 19], although the timing to be localized at the division site differs each other [20, 21]. We first treated cells expressing GFP-Bgs1 with TNM-A to find that GFP-Bgs1 was localized at sites with thick ectopic cell wall (Figure 3A). In the 1P and 2P cells, the signal of GFP-Bgs1 was observed at one cell tip and two cell tips, respectively (Figure 3B, Figure S7). In addition, in cells possessing a septum such as T1, T2, and T3 cells, GFP-Bgs1 was localized at sites with thick ectopic cell wall. The localization of Bgs1 is tightly regulated during the cell separation process and the simultaneous localization of GFP-Bgs1 at the cell tips and the septum is not observed under the normal cultivation condition: Bgs1 is localized at cell tips in the interphase and at the cell division site in the mitotic cells [12] (Figure 3B). In contrast, the simultaneous localization of GFP-Bgs1 at the cell tips and the septum was observed in T2 and T3 cells under the TNM-A challenge, suggesting that TNM-A forced polarized persistent localization of GFP-Bgs1 at the cell tips and septum (Figure S7). When cells expressing GFP-Bgs4 or Ags1-GFP were treated with TNM-A, similar localization patterns as that of GFP-Bgs1 were observed (Figure S8, S9, S10). The fluorescent signal of GFP-tagged GS proteins at the sites with thick ectopic cell wall was increased, which was more apparent at cell tips (Figure 3C). Transient increase of the fluorescent signal was observed at septa. In contrast, the fluorescent signal of GFP-tagged GS proteins at cell tips increased in a time-dependent manner. These results suggested that TNM-A induced an increased, persistent, and polarized localization of the GSs at the areas of ectopic cell wall synthesis.

**Figure 3.**
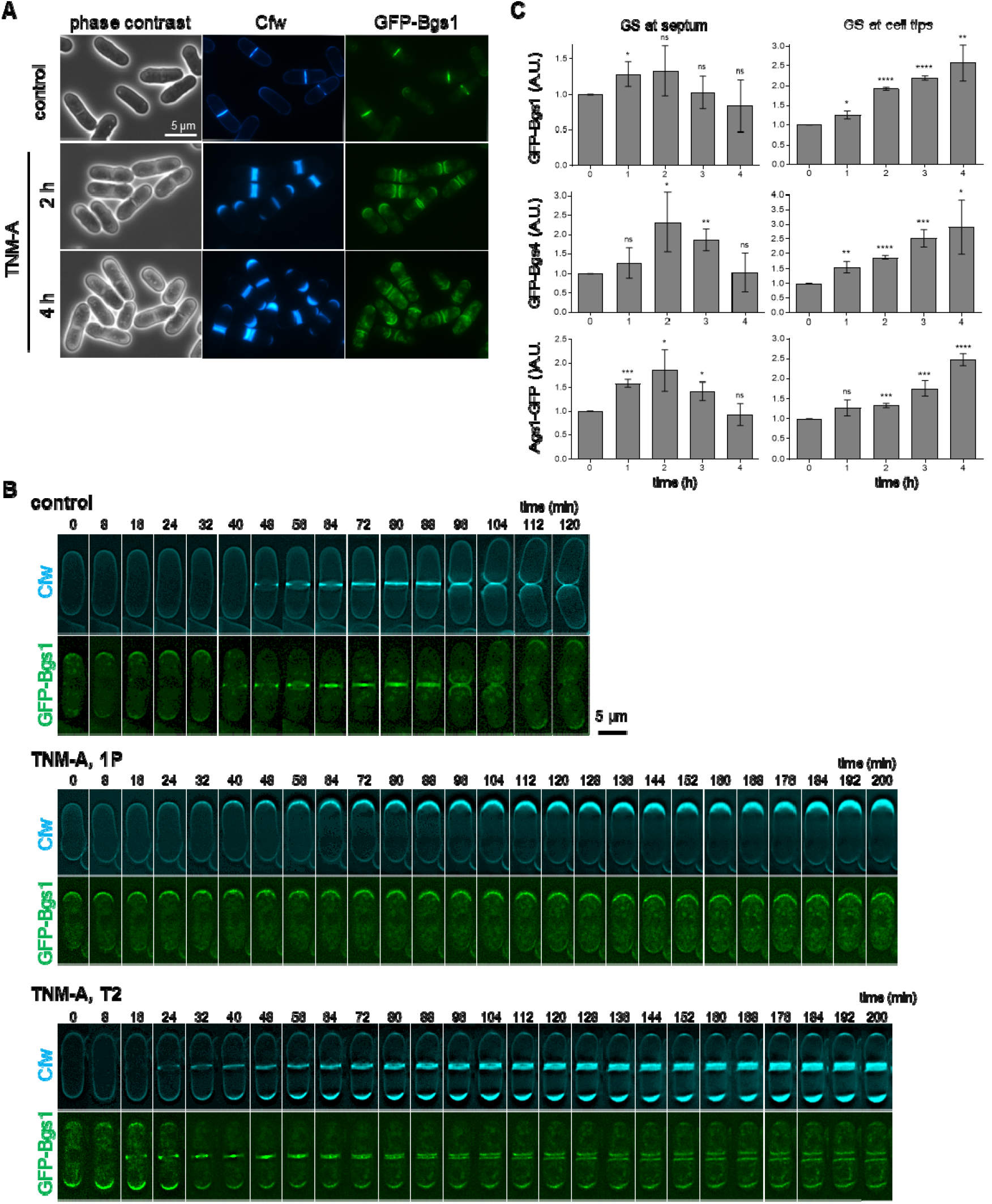
Increased, persistent localization of glucan synthases Bgs1, Bgs4, and Ags1 after TNM-A treatment. (A) Localization of GFP-Bgs1 after treatment with TNM-A (2 μg/mL). Images of phase contrast, and calcofluor white (Cfw, linear 1,3-β-glucan) and GFP-Bgs1 fluorescence microscopy are shown. (B) Time lapse images of cells expressing GFP-Bgs1 in the presence of TNM-A. GFP-Bgs1 cells were incubated in YES medium at 28 °C with TNM-A (2.0 µg/mL) and the images were captured every 8 minutes to observe 1,3-β-glucan (Cfw) and GFP-Bgs1. Top: control cell treated with DMSO. Middle and bottom: 1P-and T2-type cells with TNM-A as in figure 1C, respectively. (C) Quantification of the glucan synthases at septum and cell tips. Cells expressing GFP-Bgs1, GFP-Bgs4, or Ags1-GFP were treated with DMSO or TNM-A and imaged at the indicated times of treatment with TNM-A as in A. The fluorescence intensity of GS with TNM-A was normalized by that with DMSO. Data represent the mean ± SD (three independent experiments, 14-48 cells for septa, total 478 cells, and 45-267 cells for cell tips, total 2778 cells, were counted in each condition). Each time point with TNM-A was analysed with respect to the control with DMSO by the Student’s t-test: n.s. (not significant) p >0.05, * p <0.05; ** p <0.01, *** p <0.001, **** p <0.0001.

To examine the relationship between the GS and the cell survival in the presence of TNM-A, we tested several temperature-sensitive mutant strains of the GS (Figure S11). The mutants examined have decreased catalytic activity at semi-permissive temperature but still maintain cell growth. The *bgs1* mutant strains, including *cps1-12, cps1-N12*, and *cps1-191* strains, showed increased cell death in the presence of TNM-A. The *ags1-664* strain also showed decreased tolerance to TNM-A. In contrast, the sensitivity of the *bgs4* mutant strains, such as *cwg1-1* and *cwg1-2*, did not show increased sensitivity. These results suggest that the increased glucan synthesis, especially by Bgs1 and Ags1, protect cells against TNM-A that eventually kill cells through membrane damage [10].

### Analysis of the localizome of proteins at sites with thick ectopic cell wall in cells treated with TNM-A (Figure 4, Figure S12, Table S1)

To unveil the mechanism underlying the increased, persistent localization of GS proteins by TNM-A, we conducted a proteomic screen to identify proteins that are localized at the cell tips or septa with the thick ectopic cell wall in the presence of TNM-A. Our screen is based on the localizome catalog for the whole protein subcellular localization that was generated by observing C-terminally YFP-tagged proteins [22]. The localizome database annotated protein localization by more than one category among 19, such as nucleus, mitochondrion, ER, etc. In that catalog, 392 proteins are annotated to be localized at “periphery” and/or “cell tip and site of septum formation”, which were examined in this study (Figure 4A). Cells expressing each YFP-tagged protein were treated with TNM-A for two hours, then observed under fluorescence microscopy. After first round screen followed by second round screen to confirm the reproducibility, we identified 32 proteins that are localized at/around thick ectopic cell wall (Table S1).

**Figure 4.**
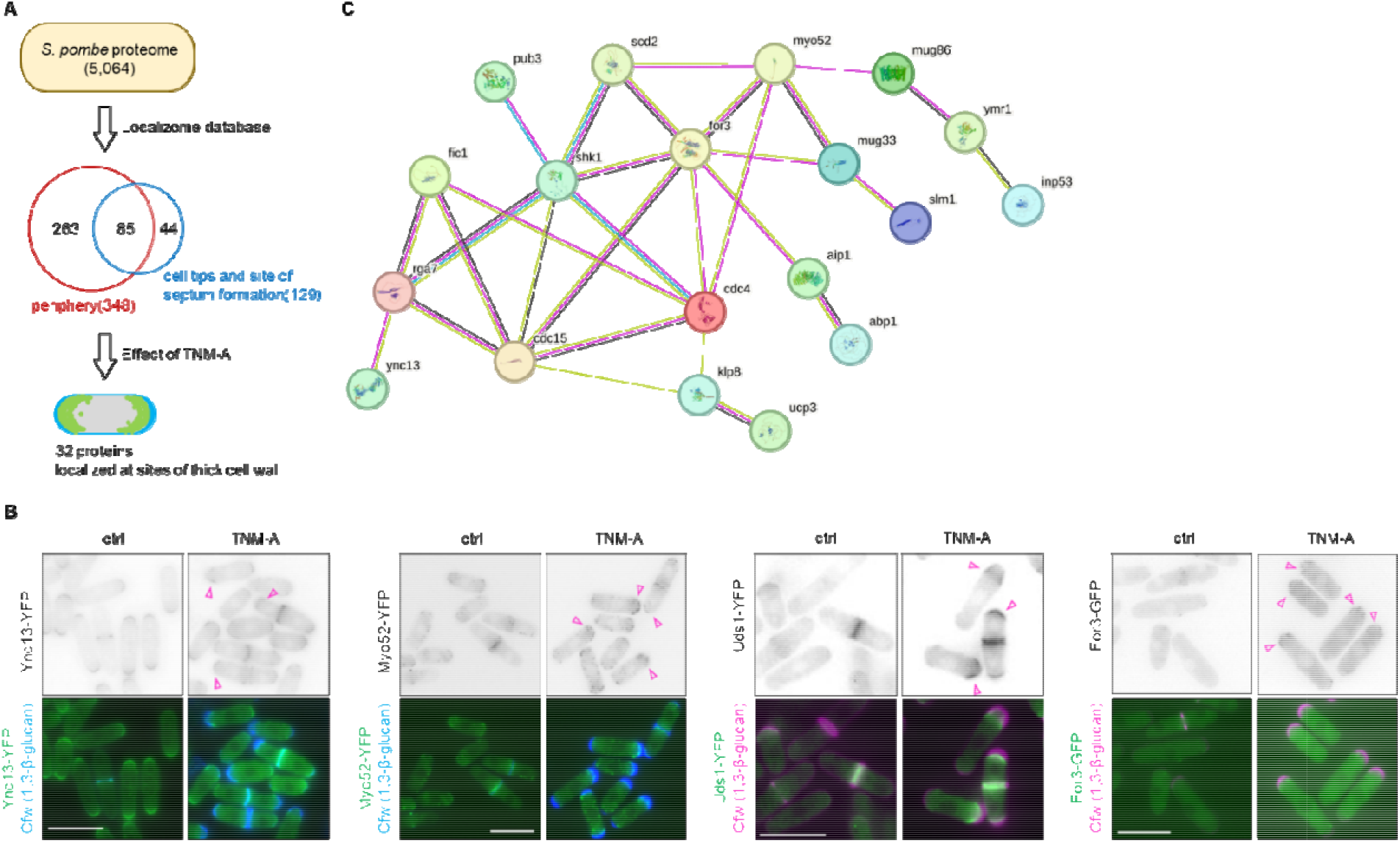
Screen for proteins involved in the action of TNM-A. A. Schematic of the screen based on the localizome database. 392 proteins among the fission yeast proteome were screened to identify 32 proteins that are localized at sites with thick ectopic cell wall. B. Examples of YFP-tagged proteins localized at polarized sites with thick ectopic cell wall. Cells were treated with TNM-A (2 μg/ml) for 2 h. The cell wall and corresponding 32 proteins were visualized by Cfw and YFP fluorescence microscopy, respectively. Scale bar, 10 μm. C. Protein-protein network drawn for 19 proteins among the 32 hit proteins. The remaining 13 were not connected by STRING.

Most proteins among the 32 hit proteins were detected as cytoplasmic dots at cell polarity sites, while some of them were observed at the cell membrane of the poles, such as Mug33, Ync13, and Uds1 (Figure 4B, Figure S10). The 32-protein information was subjected to a web-based tool STRING for visualizing the protein-protein interactions [23]. More than two thirds of the proteins showed interactions with other proteins, while one major network consisting of 19 proteins was drawn (Figure 4C). Myo52 and For3 are the hub proteins showing interactions with 10 and eight proteins, respectively. Myo52 and For3, involved in the membrane-trafficking and establishment of the cell polarity, were detected as dotted structures, at sites with thick ectopic cell wall (Figure 4B). Shk1 and Scd2 are commonly found as interactors of Myo52 and For3. Shk1, Scd2, and For3 are interactors of Cdc42, which is not included in the screen since the C-terminus should be processed for the correct localization of Cdc42. Cdc15, Abp1, and Fic1 that have seven, six, and five interactors, respectively, are the proteins that are localized at cell polarity sites. These results imply that TNM-A activates the cell polarity machinery, i.e. Cdc42, by which excess GS proteins might be exported to the cell tips and septa.

### Activation of Cdc42 by TNM-A (Figure 5)

Cdc42 is a widely conserved small GTPase, regulating polarized cell growth and cell morphogenesis. Localization of active Cdc42-GTP can be visualized by expressing the GFP-tagged Cdc42/Rac-interacting binding (CRIB) domain [24]. The fluorescent signal of CRIB-GFP is detected at cell tips at the interphase or at the site of septation under normal culture conditions (Figure 5A). In the presence of TNM-A, the signal of CRIB-GFP at cell tips became more intense in a time dependent manner, which was supported by line scan analyses (Figure 5B). As Cdc42 is involved in the polarized growth, TNM-A might increase the transport of GSs to the cell-polarity sites through activation of Cdc42.

**Figure 5.**
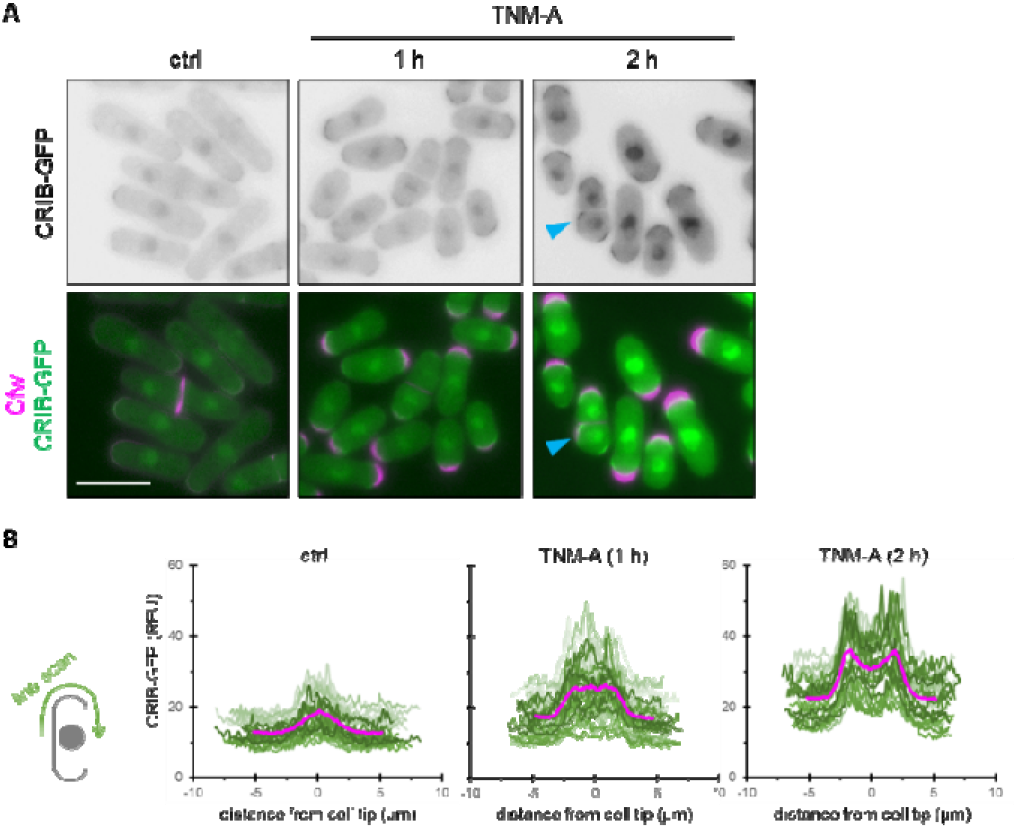
Activation of Cdc42 by TNM-A. (A) Polarized localization of CRIB-GFP. Cells expressing CRIB-GFP were treated with TNM-A (2 μg/ml) and stained with Cfw (magenta). The arrowhead (blue) indicates the ectopic activation of Cdc42 and deposition of the cell wall material. Cells were visualized by Cfw, and GFP fluorescence microscopy. CRIB, Cdc42/Rac-interacting binding (probe for active Cdc42-GTP). Scale bar, 10 μm. (B) Quantitation of the CRIB-GFP fluorescence. The intensity of the fluorescence was traced along the cell tips as shown in the schematic figure to the left. The thick magenta line is the average of line scans from 30 cells in three independent experiments.

### Not only transport along actin cables, but also random walk of the vesicles is up-regulated by TNM-A (Figure S15-16)

We investigated if the membrane transport along actin cables is required for the action of TNM-A. Mug33, one of the 32 proteins that were detected at sites with thick ectopic cell wall (Figure S12), is localized at the plasma membrane in a polarized manner, which depends on For3 and Myo52 [25]. Notably, both For3 and Myo52 were detected at sites with the thick ectopic cell wall (Figure 4B). In the absence of For3 or Myo52, the forced accumulation of Mug33 by TNM-A was rarely observed (Figure S13). Instead, intense cytosolic fluorescent signals of Mug33-YFP were observed. These results indicated that TNM-A can upregulate the membrane trafficking along the actin cables.

We next tested if the transport along the actin cables is required for the persistent localization of GS with TNM-A. In the *for3*Δ or *myo52*Δ cells, GS proteins including Bgs1, Bgs4, and Ags1 were still observed at the site of septation (Figure 6, S14). However, their signals at cell tips were weak or rarely observed and dispersed in the cytoplasm, indicating that the polarized transport of GS proteins to the growing tips is decreased or the localization at the growing tips is unstable in the absence of the actin-cable dependent transport. In contrast, the cell-tip localization of GS proteins was observed after TNM-A treatment, which was comparable, or even much higher, to that of the control cells (Figure 6, S14). The characteristic short, oval cell morphology of *for3*Δ or *myo52*Δ cells was not restored by the addition of TNM-A. These data revealed that the vesicle transport along actin cables is not requisite for the increased, persistent localization of GS proteins by TNM-A and suggests that the random walk of exocytic vesicles to the cell polarity sites is activated by TNM-A [26].

**Figure 6.**
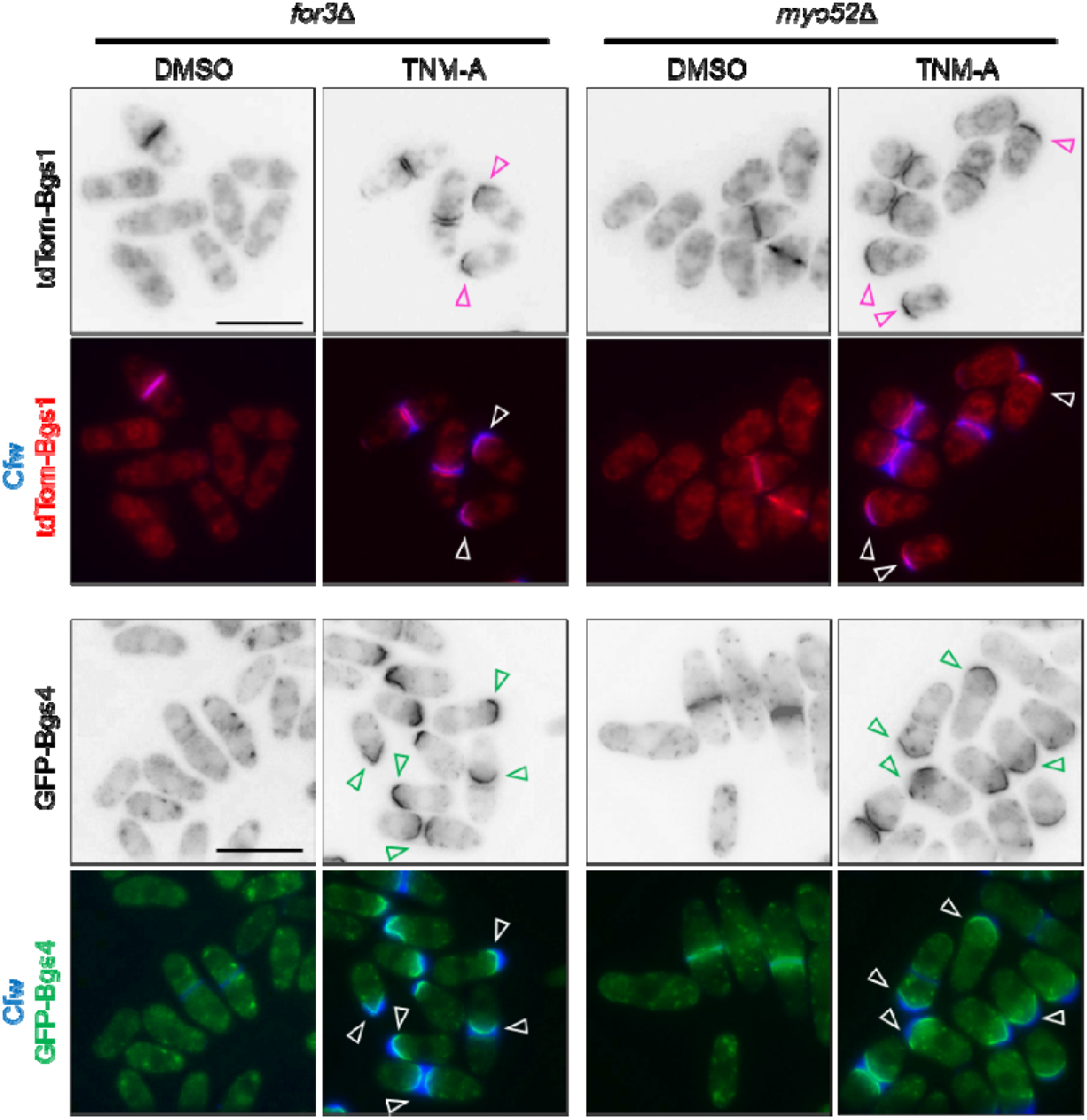
Effect of for3 or myo52 deletion on the localization of Bgs1 and Bgs4. *for3*Δ or *myo52*Δ cells containing tdTom-Bgs1 or GFP-Bgs4 were treated with DMSO (control) or TNM-A (2 μg/ml) for 2 h. Cells were stained by Cfw (mainly linear 1,3-β-glucan) and observed under phase contrast and Cfw, tdTom, and GFP fluorescence microscopy. Scale bar, 10 μm. Open yellow triangle; accumulation of tdTom-Bgs1 and GFP-Bgs4 at the poles.

## DISCUSSION

Sterols have distinctive chemical properties among the membrane lipids: the polar function of the sterols is a hydroxy group, being smaller than other membrane lipids such as phospholipids, and the hydrophobic portion has a rigid, planar structure. This unique feature enables tight interactions with lipids with long, saturated alkyl chains, such as sphingolipids, by which sterols can exhibit multiple biological functions. This study demonstrates that capturing sterols in fission yeast by TNM-A activates Cdc42, induces increased, persistent localization of GSs at cell-polarity sites, and leads to overproduction of glucan, which suggests that plasma membrane sterols function to maintain proper cell polarity and cell wall biosynthesis.

The thick ectopic cell wall phenotype has been reported for several mutant strains. In contrast to TNM-A, genetic mutations often produce round cell morphology with thick cell wall, indicating the loss of the cell polarity. For example, cells with decreased activity of Ags1 [27], dysregulation of the Bgs1 localization by depletion of Sbg1 that is an associate protein of Bgs1 [28], and expression of constitutively active Rho1 have thick cell wall and lose the cell polarity [11, 29]. Instead, the thick cell-wall phenotype of TNM-A-induced cells seems to be similar to the phenotype of *css1* mutant strains [30, 31]. Css1 is a plasma membrane inositol phosphosphingolipid phospholipase C, whose dysfunction leads to increase in the cellular amount of mannosylinositol phosphorylceramide (MIPC) and reduction in the membrane fluidity of inner leaflet of the plasma membrane. TNM-A might also modulate the membrane fluidity of the plasma membrane. In fact, TNM-A induces phase separation in artificial lipid membranes and modulates membrane order in cultured mammalian cells [6]. Curiously, similar thick cell wall phenotype was observed with other antifungal compounds, such as heronamides and tubimoside I [32, 33]. Heronamides bind to phospholipids possessing long alkyl chains. Although the cellular target of tubimoside I is missing, it can also target the cell membrane as many saponins are known to perturb the cell-membrane integrity. Taken together, the dysregulation of the membrane properties, especially the fluidity, seems to induce cell wall overproduction without losing cell polarity.

1,3-β-glucan and α-glucan are the two major components of the thick ectopic cell wall that is formed by the action of TNM-A (Figure 2, Figure S4). The increase of the two components is consistent with the increased, presistent localization of GS (Figure 3, Figure S7-S10). The three GS proteins, Bgs1, Bgs4, and Ags1, showed polarized, sustained localization at the cell tips and septa in the presence of TNM-A. In particular, it is characteristic that GSs are localized simultaneously to cell tips and septation sites in TNM-A-treated cells, whereas GSs in fission yeast are regulated by membrane trafficking in synchrony with the cell cycle and do not localize simultaneously to cell tips and septation sites [12, 18-20]. The increase of the active Cdc42 by TNM-A may contribute to the increased GS trafficking to the plasma membrane. It is also assumed that internalization of GS from the plasma membrane might be decreased. It would not be surprising if GS internalization is decreased, since the membrane environment is expected to be altered by TNM-A as described above. This mechanism might be shared with the *css1* mutants [31]. Although the time-course analyses of GS proteins were not reported, GS proteins were shown to be localized adjacent to the thick cell wall at least after 3 h incubation at the restricted temperature of the *css1* mutant.

In mammalian cells, cholesterol depletion was reported to activate Cdc42, which leads to activation of the downstream events such as membrane trafficking and filopodia formation [34-38]. In fission yeast, the active Cdc42 was detected just beneath the thick ectopic cell wall, which was increased by TNM-A in a time-dependent manner (Figure 5). The increased active Cdc42 likely upregulates the trafficking of GSs to the polarity sites. Furthermore, it might also contribute to the polarized cell shape that was kept while synthesizing excess cell wall. Aside from whether the molecular mechanisms are conserved between yeast and humans, it will be interesting to look at how the activity of Cdc42 and the cell polarity are regulated by the membrane sterols.

Comprehensive analysis is often effective to narrow down the candidate molecules that function in the phenomena of interest. In this study, localizome screen implied that TNM-A may activate Cdc42. In contrast to the growth-inhibition-based screen, such as chemical genomic analyses [39, 40], localizome screen can furnish unique results, e.g. proteins that are involved in the mechanism of action (MoA) of a compound but do not affect the cell growth, can be identified [22]. Even in the case of compounds that do not show toxicity but affect the cellular status, the localizome screen would be effective. On the other hand, the ORFeome strains used in this study are expressing proteins tagged at the C-terminus, so correct localization information cannot be obtained for proteins such as small GTPases and GS. When tagged at the C-terminus, small GTPases such as Cdc42 and Rho family are diffused in the cytoplasm, while none of the GS are functional and delivered to the plasma membrane but are found in the ER or form cytoplasmic dots. In order to conduct comprehensive analyses of the subcellular localization of proteins, it is necessary to develop a strain collection consisting of cells that express functional, labelled proteins.

In this study, we demonstrate that temperature-sensitive mutants of *bgs1* and *ags1* showed hypersensitivity to TNM-A, which suggests that glucan overproduction is one of the stress responses to protect cells from the membrane damage by antifungals (Figure S11). Although we demonstrated the activation of Cdc42 and abnormal localization of GSs by TNM-A, the molecular mechanism underlying this stress response largely remains to be elucidated. As TNM-A generates thick ectopic cell wall gradually and the phenotype is drastic, this molecule will be a useful chemical biology tool to elucidate the cellular systems sensing the cell membrane perturbation.

### Limitations of the study

Cdc42 was shown to be activated by TNM-A. However, we could not investigate the involvement of Cdc42 by genetic or chemical genetic inactivation, since TNM-A requires the Cdc42 activity to bind to the cell. Inactivation of Cdc42 followed by attenuation of exocytosis decreases the amount of ergosterol in the plasma membrane, which is the target molecule of TNM-A [41]. For this purpose, a new technique by which Cdc42 can be quickly inactivated just after cellular binding of TNM-A is requisite. Another point to be tested is the behaviour of membrane proteins other than GSs. Which membrane proteins in addition to GSs are accumulated at the PM? TNM-A would be able to classify membrane proteins: membrane proteins that are accumulated at cell polarity sites by TNM-A, and those that are not. The difference might be due to the difference of the transport system, the size of the proteins, or unexpected properties. Finally, Rho1 was previously shown to be required for the thick cell wall phenotype of TNM [10]. Analyses of the involvement of Rho1 in the activation of Cdc42, increased, persistent localization of GSs, and thick cell wall will unveil novel interactions between key small GTPases Cdc42 and Rho1.

## METHODS

### Yeast strains and cultivation

Yeast strains used in this study are listed in Tables S2. Gene deletion mutants of *for3* and *myo52* were generated by random sporulation using the Bioneer library v5.0 [42]. Correct gene deletion was confirmed by colony PCR. Cells expressing C-terminally YFP-tagged proteins were prepared previously [22]. Yeast cells were cultivated in Edinburgh minimal medium 2 (EMM) or YES. YES is a rich medium composed of yeast extract, glucose, and 225 mg/L each of five supplements (Ade, Ura, L-Leu, L-His, and L-Lys) [43].

### Screening of ORFeome strains

Strains cultivated on YES plates were inoculated into EMM and cultivated at 28 °C overnight. The cell culture was diluted to 0.2 OD_595_, followed by further cultivation for 2 h. Cells were collected by centrifugation (2,500 rpm, 1 min) and resuspended in EMM supplemented with Cfw (10 μg/ml) (0.35-0.40 OD_595_). A portion of the cell suspension (150 μl/well) was poured into 96 well plates that were coated by lectin and left for 5 min [44]. Cells not attached to the bottom were washed out twice by MM+Cfw (120 μl/well). Wells were filled with MM+Cfw containing DMSO (0.2 % v/v) or TNM-A (2 μg/ml) (120 μl/well), incubated for 2 h, and observed under fluorescence microscopy. After the first screen, the reproducibility was confirmed in the second screen to identify 32 proteins that are localized at sites with thick ectopic cell wall (Table S1).

### Fluorescence microscopy

Images were obtained with two systems: KEYENCE BZ-X710 fluorescence microscope equipped with a PlanApo 100× lens and BZ-X viewer software; a Leica DM RXA fluorescence microscope equipped with PL APO 63×/1.32 oil PH3 and PL FLUOTAR 100x/1.30 oil PH3 objectives, a digital camera (DFC350FX; Leica Biosystems), and CW4000 cytoFISH software (Leica Biosystems), and processed (brightness, contrast, and/or sharpness) with Adobe Photoshop CS2 software. Time-lapse video imaging was performed essentially as previously described [19, 44]. For cell wall staining, early or mid log-phase cells were cultivated at the corresponding temperature in YES or EMM liquid media. The cell wall was visualized by adding a solution of Calcofluor White (Cfw; 10 or 50 μg/ml final concentration, Fluorescent Brightener 28, Sigma-Aldrich) to the sample and using the appropriate filter. Cells with GFP-, YFP-, or RFP-labelled proteins were directly visualized using appropriate filters. All the analyses were repeated at least three independent experiments, and representative images were selected to be shown. Graphical data were calculated from at least three independent experiments. Line scan analyses were conducted by ImageJ software. The numbers of experiments and of total septa or cells analyzed are shown in each case.

Concanavalin A conjugated to fluorescein isothiocyanate (FITC) was used to indirectly quantify the galactomannoproteins of the outer layer of the cell wall as previously described [45]. Equal amounts of log-phase cells with and without the nuclear marker Hht1-RFP, and cultivated or not in the presence of TNM-A, were mixed and analyzed. Fluorescence intensity was quantified by ImageJ software. In each experiment, at least 6 images were captured for each 2-h interval over a 6-h period of growth in the presence of TNM-A. In all cases, the background of the photo was subtracted using a fixed ROI.

For GFP fluorescence intensity analysis of the GS GFP-Bgs1, GFP-Bgs4, and Ags1-GFP fluorescence in the septum membrane, equal amounts of log-phase cells either in the absence or presence of TNM-A, and with or without Hht1-RFP, were mixed and analyzed. Cfw staining of cells growing in the presence of TNM-A is very different from untreated control cells. Therefore, combinations of cells of the same strain incubated either in the absence or presence of TNM-A, were also directly used. A rectangle covering the entire septum or cell pole area was drawn in each cell. The rectangle area (ROI) was always the same. Fluorescence was compared using the ‘RawIntDen’ value. In each image, the average ‘RawIntDen’ value of the septa or cell poles of each strain was obtained. In each experiment, at least 6 images were captured for each 2-h interval over a 6-h period of growth in the presence of TNM-A. In all cases, the background of the photo was subtracted using a fixed ROI.

### Electron microscopy

Wild-type cells in mid-log phase were cultivated with DMSO (0.2%) or TNM-A (2 μg/ml) for 2 h at 27 °C. Cells were washed by PBS (100 mM phosphate buffer, pH 7.2, 150 mM NaCl) and fixed with 2% glutaraldehyde in PBS for 2 h at 4 °C. Fixed cells were washed with PBS and stored at 4 °C before electron microscopy. The specimens were treated as described before [46] and observed by a scanning electron microscope SU9000 (Hitachi High Technologies, Tokyo) operating at 0.8 kV, and a transmission electron microscope JEM-1011 (JEOL, Tokyo) operating at 100 kV.

### Labeling and fractionation of cell wall polysaccharides

Labeling the cell wall with D-[U-^14^C]-glucose and the fractionation into α-glucan, β(1,3)-glucan, β(1,6)-glucan, and galactomannoproteins were performed as described previously [47, 48] with some modifications [49]. In this study, cells were labeled with D-[U-^14^C]-glucose (3 µCi/ml) for 6 h prior addition of TNM-A (2.0 μg/ml) and collected at the indicated times with TNM-A. For the analysis of cell wall polysaccharides synthesized at different times with TNM-A, cells were pulse-chase labeled with D-[U-14C]-glucose (8 µCi/ml) for 2 h in the different intervals with TNM-A (0 to 2 h, 2 to 4 h, and 4 to 6h). All experiments were performed in duplicate and values were calculated from at least three independent cell cultures.

### 1,3-β-glucan synthase assay

The activity of the glucan synthase (GS) enzyme complex was analyzed as described previously [50-52]. For the GS assay with *in vitro-*added TNM-A, different concentrations of TNM-A (final concentrations: 0.05, 0.1, 0.2, 0.5, 1, 2, 5, and 10 µg/ml), or DMSO as the control, were added to the reaction mixture. For the GS assay with *in vivo-*added TNM-A, the cells were cultivated in the presence of 2.0 µg/ml of TNM-A for 0, 1, 2, and 4 h, and the enzyme extracts were obtained and assayed without TNM-A in the reaction. All GS reactions were carried out in duplicate, and the values for each strain or condition were calculated from at least three independent cell cultures.

## ACKNOWLEDGEMENTS

We are very grateful to P. Pérez for thoughtful comments, useful suggestions, and ideas. We appreciate the support that Y. Namase provided for the EM observation. We thank K.L. Gould, K. Shiozaki, P. Munz, J. Cooper, and C. Shimoda for generous gifts of strains. Vanessa S. D. Carvalho acknowledges the financial support received through a contract from Junta de Castilla y León (JCYL, Spain) and the European Regional Developmental Fund (ERDF, EU). This work was partly supported by a Grant-in-Aid for Scientific Research from the Japan Society for the Promotion of Science (25702048, 18K06717, 21H02128 to S.N.; 19H05640 and 25K24592 to S.N. and M.Y.; 23H04882 to H.K., Y.Y. and M.Y.; 17H06401 to S.N. and H.K.; 24H00493 to H.K.), PGC2018-098924-B-100 and PID2021-124971NB-I00 (Ministerio de Ciencia, Innovación y Universidades, MICIU / AEI / 10.13039/501100011033, Spain and ERDF A way of making Europe, EU); and CSI150P20 and “Escalera de Excelencia” CLU-2017-03 (Junta de Castilla y León, Spain and ERDF, EU). English editing was partly done with the aid of ChatGPT.

