## Supplementary Figure S1-S11, S13-S14, and Table S1 for "Ectopic overproduction of cell wall glucan through membrane perturbation by an antifungal peptide theonellamide A in fission yeast"

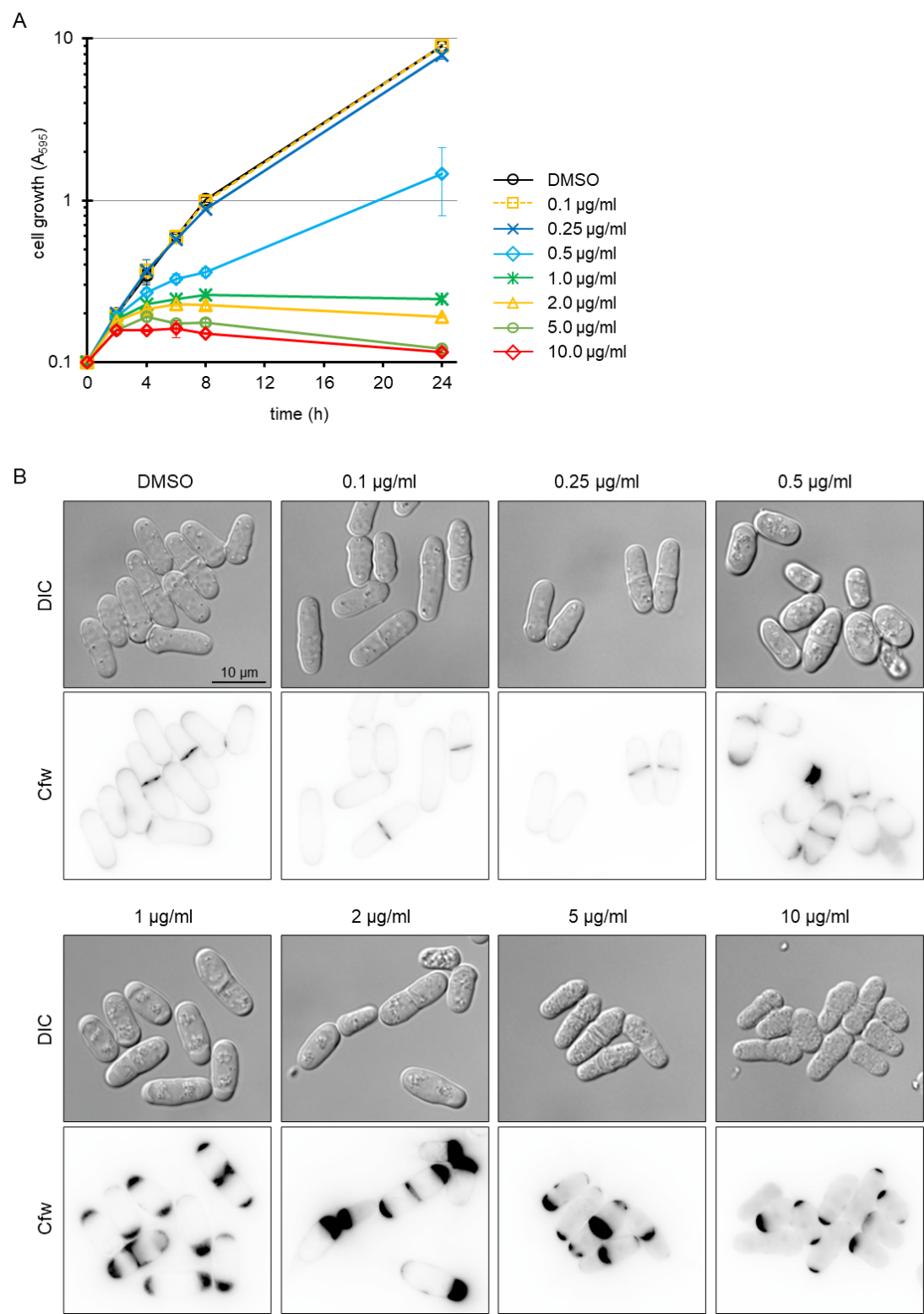

### **Figure S1. Concentration dependence of the effect of TNM-A.**

(A) Early log-phase wild-type cells growing in YES medium at 27 ºC were treated with TNM-A at various concentrations and the cell growth at A_600_ was monitored for 24 h. Data represent the mean ± SD (n = 3).

(B) Cells were treated for 7 h with increasing concentrations of TNM-A, stained with calcofluor white (Cfw), and observed under Differential Interference Contrast (DIC) and fluorescence microscopy.

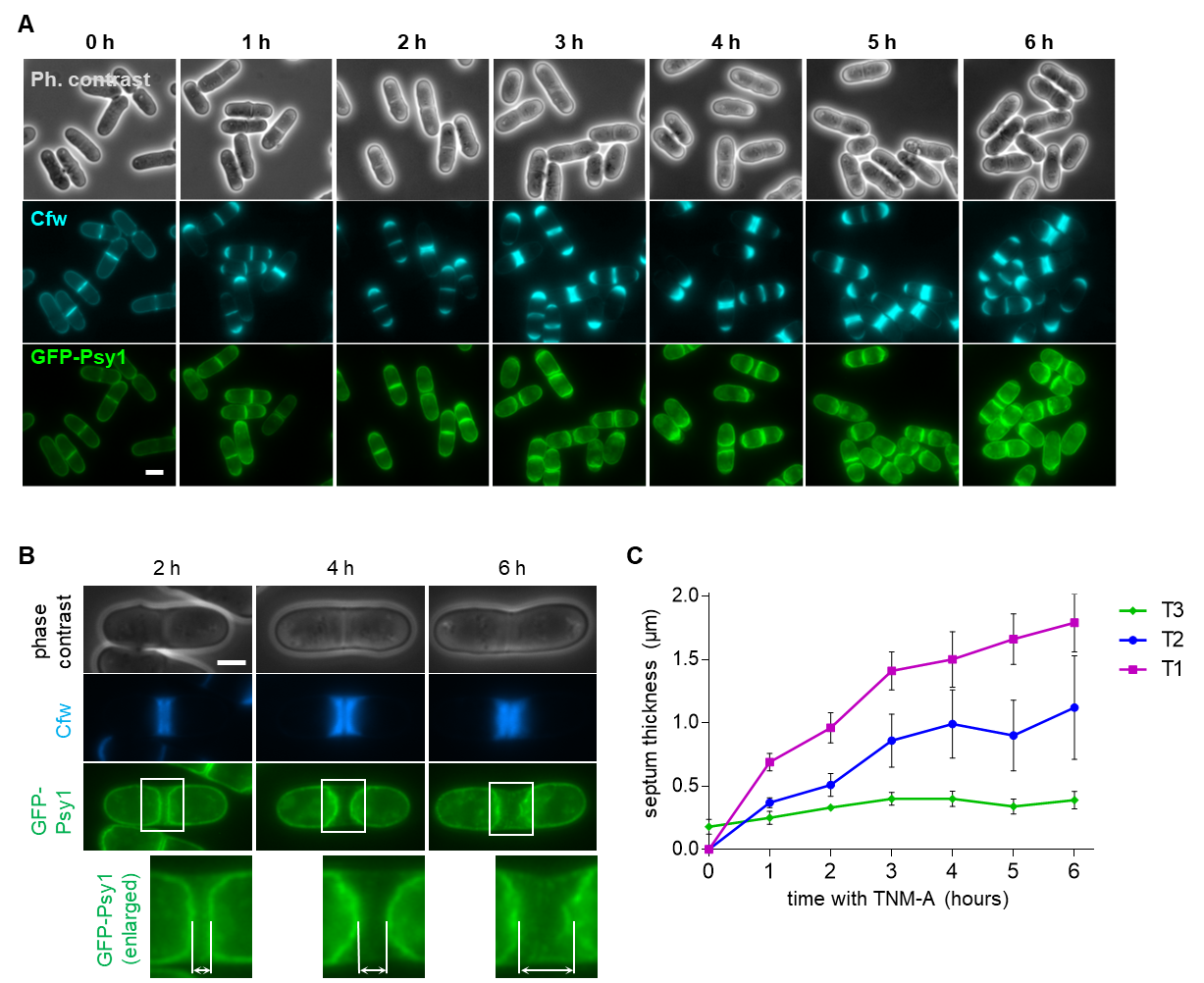

### **Figure S2. Time course analyses of the morphological changes induced by TNM-A.**

(A) Cells with GFP-Psy1 were incubated in YES medium containing 2.0 µg/mL of TNM-A under shaking at 28 °C for 6 h. At each hour, cells were collected and analyzed by phase contrast and Cfw and GFP-Psy1 fluorescence microscopy.

(B) Cells with the T1 phenotype, as in Figure 1B and C, at 2 h, 4 h and 6 h of TNM-A treatment. The septum areas with boxes are enlarged in the bottom panels and the distance between the opposing membranes surrounding the septum was measured. Scale bars, 5 μm (A), and 2.5 μm (B).

(C) Septum thickness of each of the three types of septation phenotype, T1, T2, and T3 as in Figure 1B and C, in cells growing with TNM-A for 6 h. Data represent the mean ± SD (n = 3).

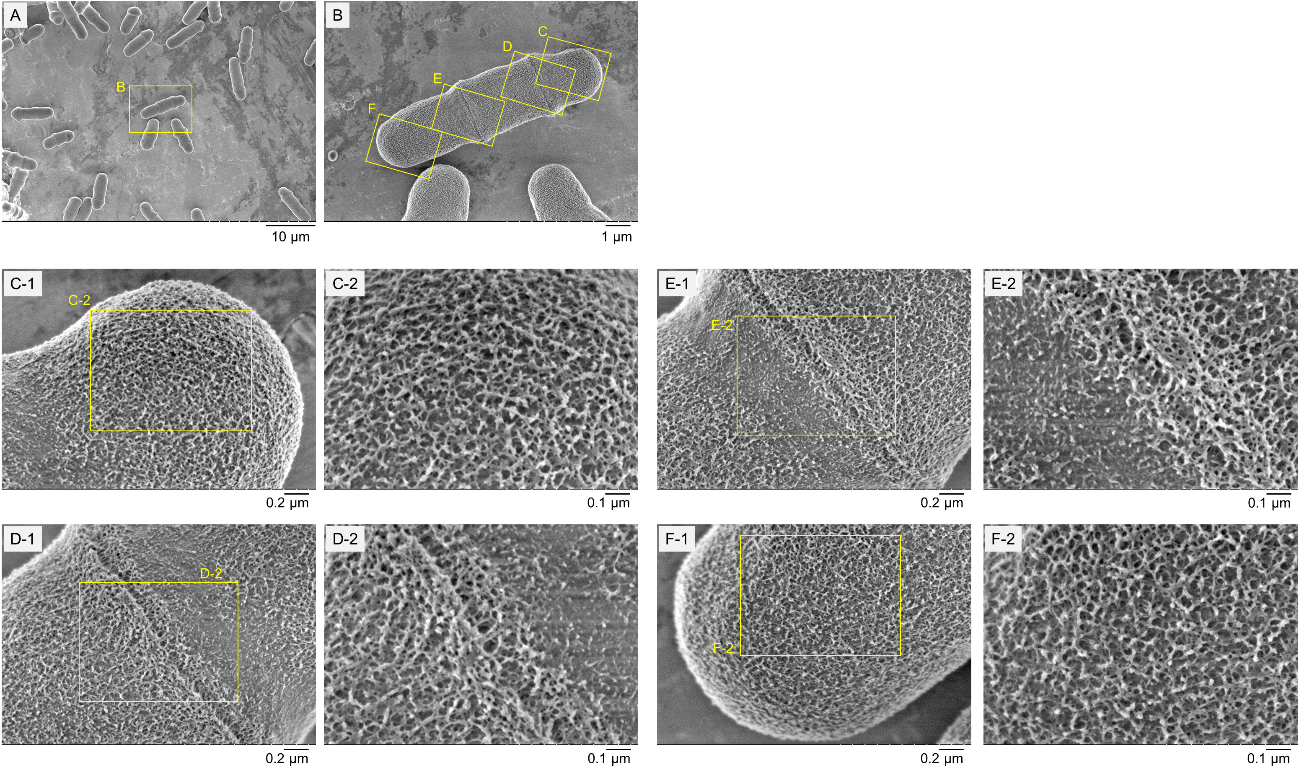

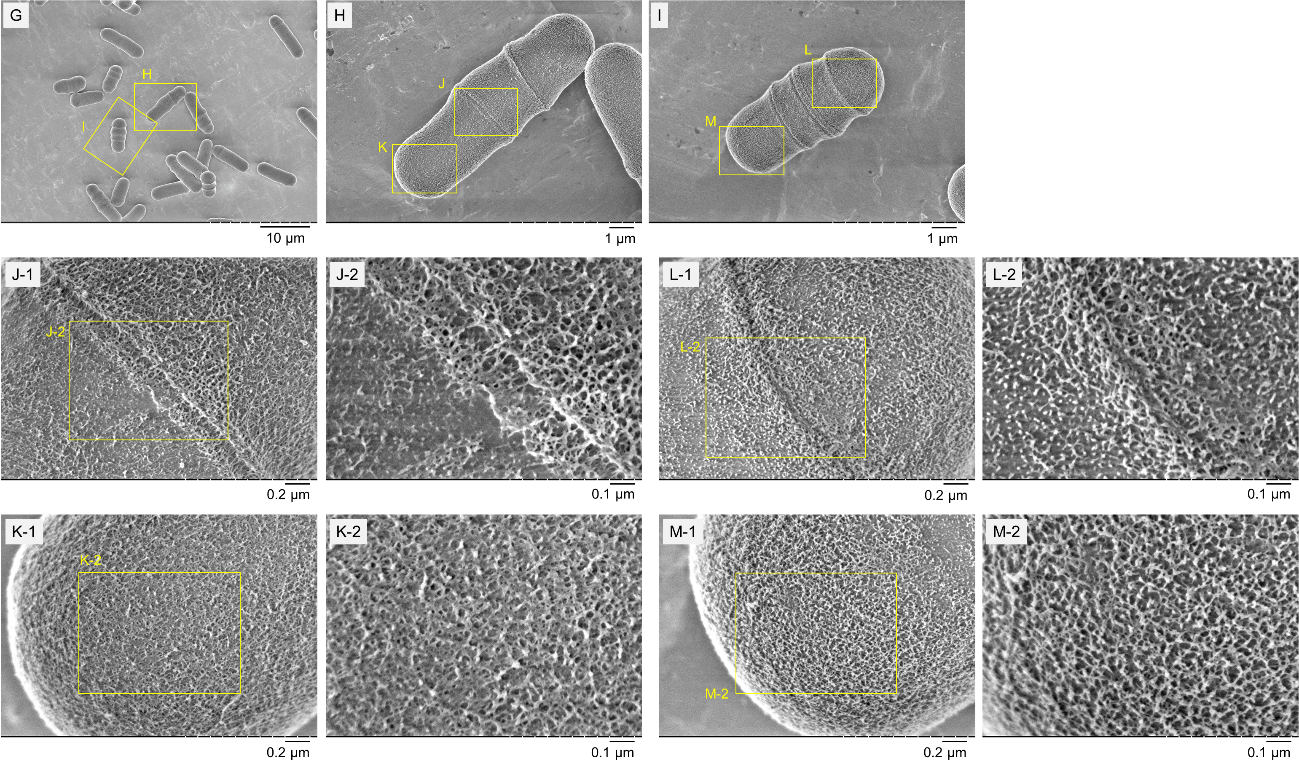

### **Figure S3. Scanning electron microscopy analyses of the cell wall surface of fission yeast cells treated with TNM-A.**

(A-F) Scanning electron microscopy (SEM) images of cells treated with DMSO. (J-M) SEM images of cells treated with TNM-A. Cells were treated with DMSO (0.5%) or TNM-A (2 μg/ml) for 2 h at 27 ˚C. Areas with boxes are marked with a letter and enlarged in the following panel with the same letter. The last areas with boxes of the most magnified images are enlarged in the right panels.

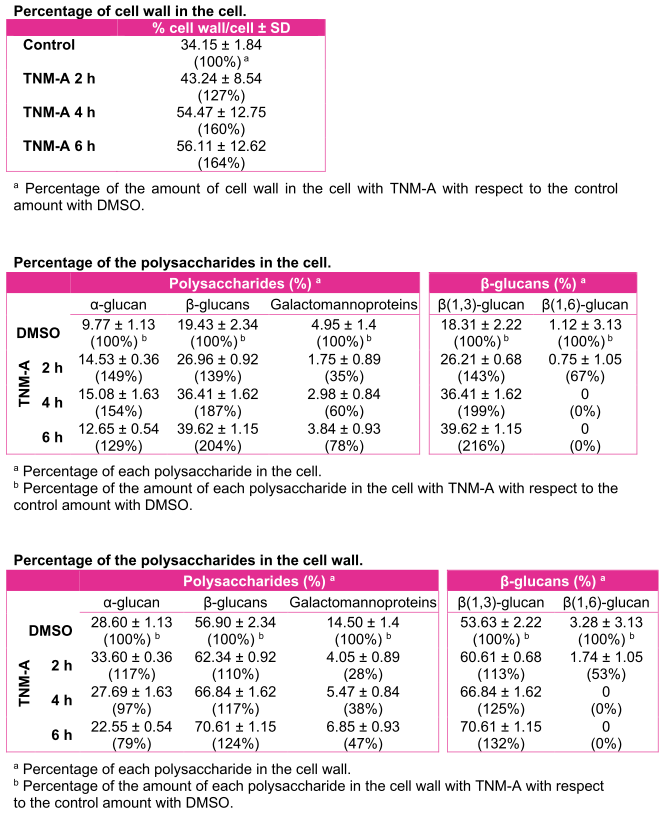

**Figure S4. The effect of TNM-A on the percentage of the cell wall in the cell, and on the percentage of the polysaccharides in the cell.**

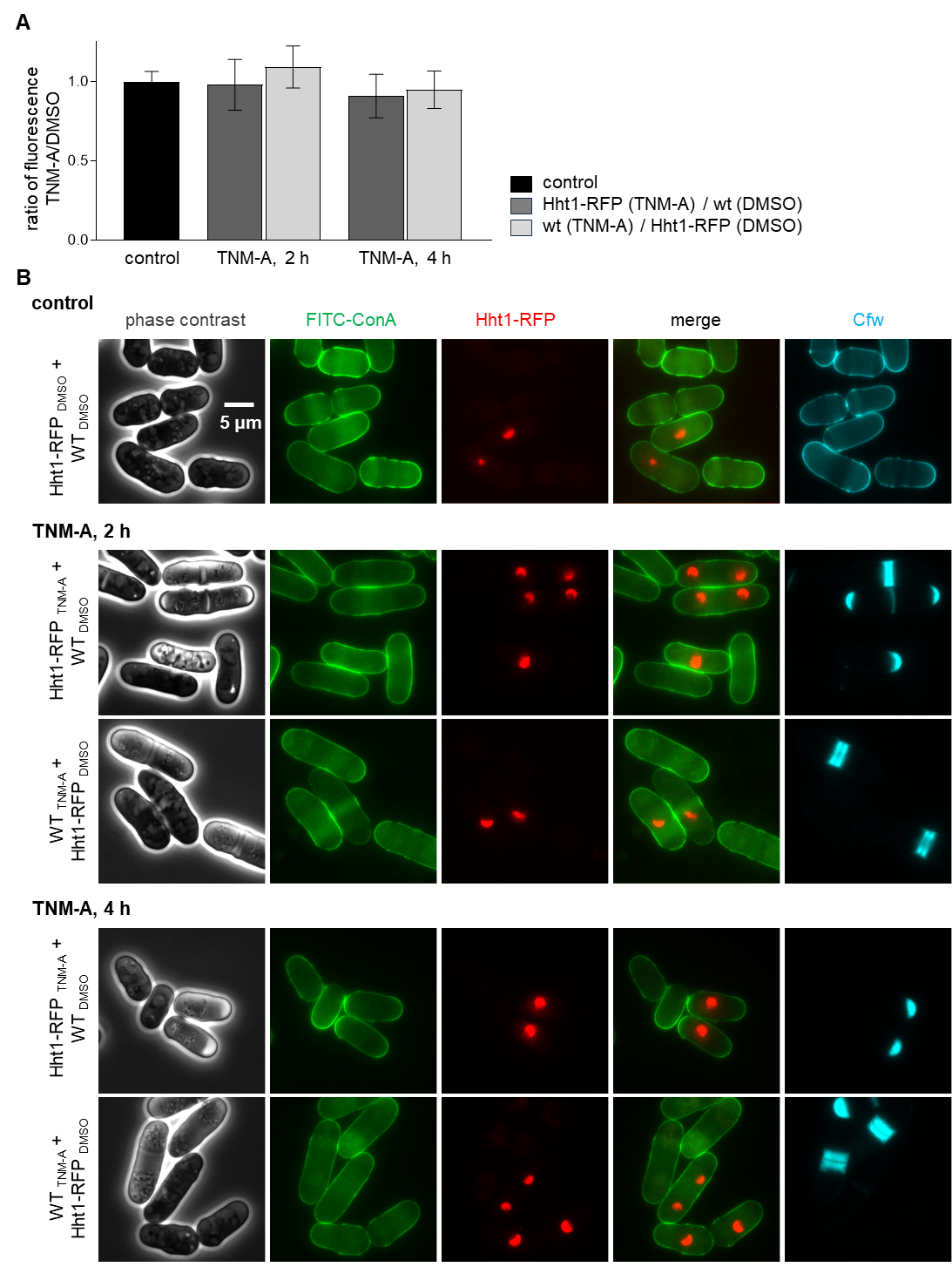

**Figure S5. Effect of TNM-A on the amount of galactomannoproteins of the outer surface of the cell wall.**

(A) Quantification of the amount of galactomannan on the cell surface. Bars represent the mean ± SD of the rate of the maximum fluorescence of the ROI on the surface of two compared strains with and without Hht1-RFP to allow their discrimination. 819 cells with TNM-A and 1650 cells with DMSO were measured.

(B) Images of the cell mixture. Early log-phase wild-type (WT) cells were treated with TNM-A, and the Hht1-RFP strain was treated with DMSO, and the opposite WT with DMSO and the Hht1-RFP strain with TNM-A, which were mixed and mounted.

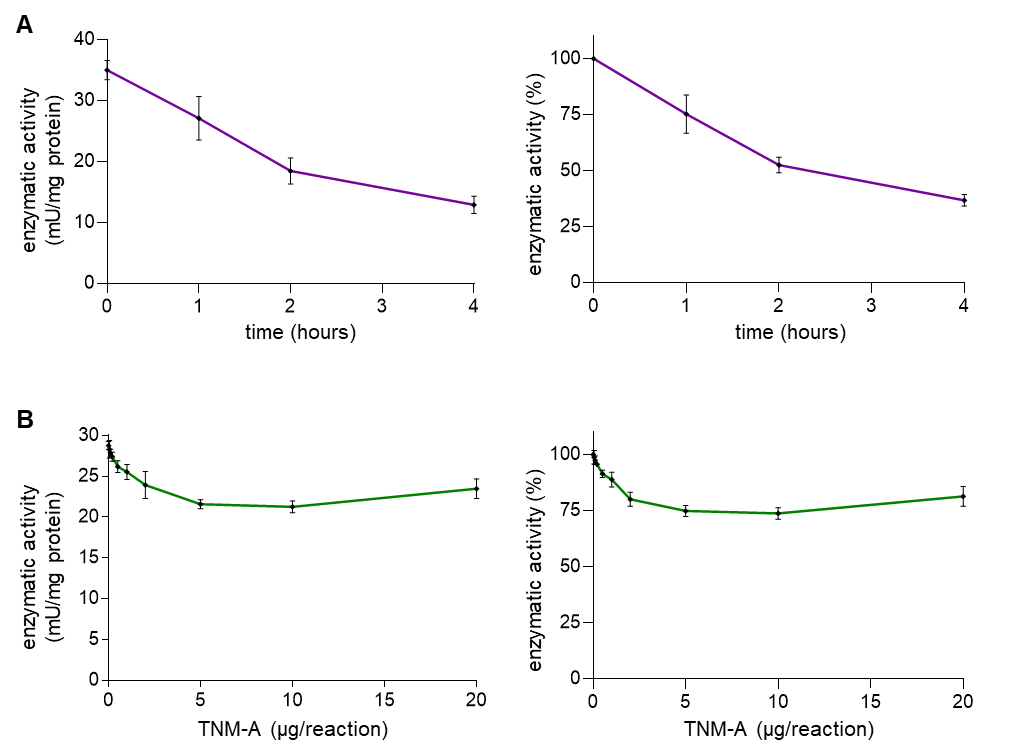

**Figure S6. The effect of TNM-A on the *in vivo* and *in vitro* 1,3-β-glucan synthase activities.**

(A) The effect of TNM-A on the cellular (*in vivo*) activity of 1,3-β-glucan synthase (GS). Early log-phase cells were cultured at 28 °C in YES with 2 µg/mL TNM-A for 0, 1, 2, and 4 hours. All cell cultures were harvested at 1.0 A_600_ (1x10^7^ cells/mL) and their membranes were extracted to quantify the GS activity.

(B) The effect of TNM-A on the *in vitro* activity of GS. Early log-phase cells were cultured at 28 °C in YES, and cell cultures were harvested at 1.0 A_600_ (1x10^7^ cells/mL) to prepare a cell membrane fraction that contains the enzyme complex. TNM-A was added to the reaction mixture to assess the effect of the compound on the GS activity.

Left graphs: specific activity is expressed as milliunits per milligram of protein (mU/mg). Right graphs: percentage values of specific activity compared to control activity in the absence of TNM-A. Data represent the mean ± SD (n = 4).

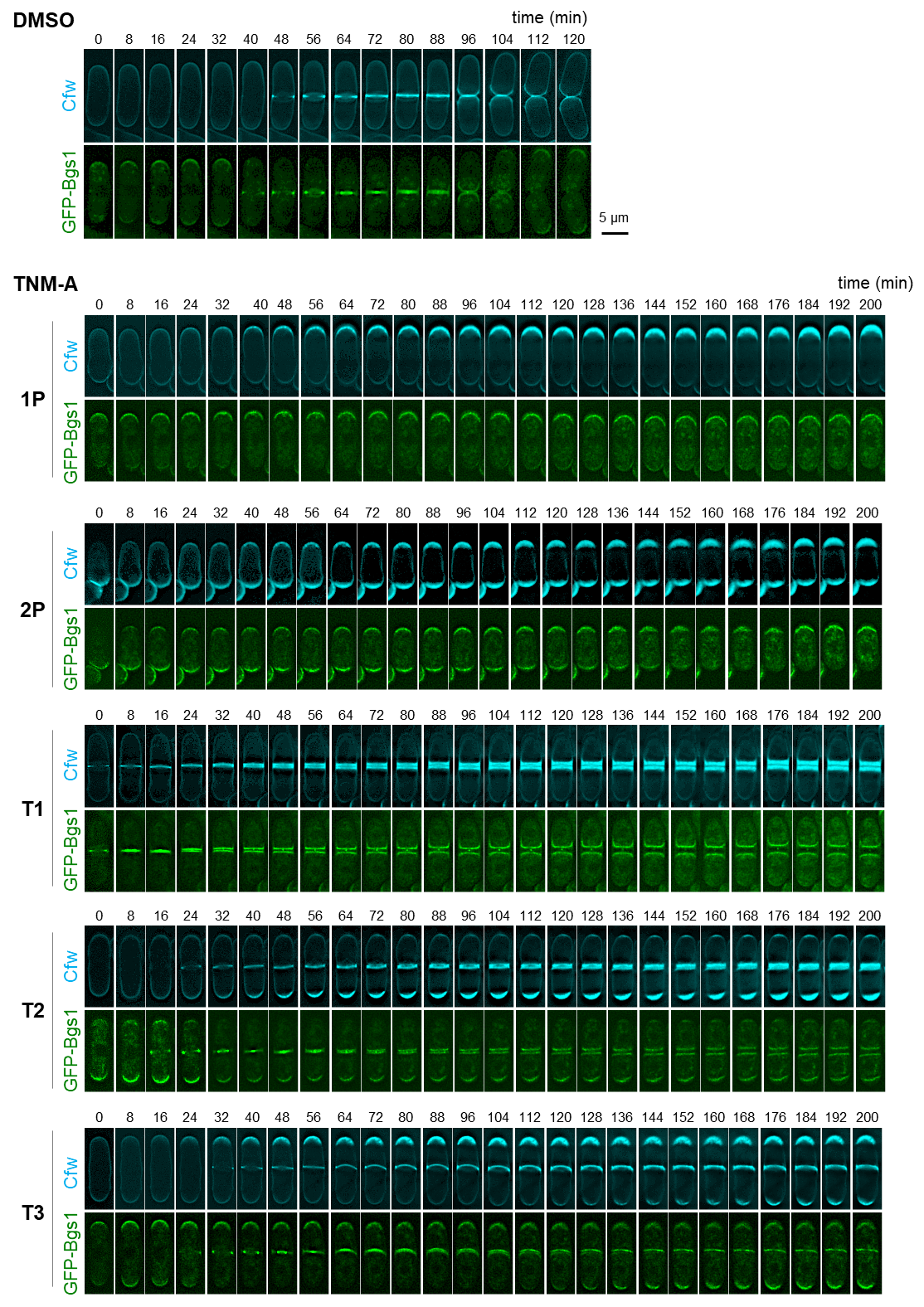

**Figure S7. Time lapse images of GFP-Bgs1 in the presence of TNM-A.**

Time lapse images of cells containing GFP-Bgs1. Cells were incubated in YES medium at 28 °C with DMSO (control) or TNM-A (2.0 µg/mL). Image capture was performed every 8 minutes for at least 120 minutes in different channels to observe Cfw staining (top, cyan) and GFP (bottom, green) fluorescence. Control cells and TNM-A treated cells with 1P, 2P, T1, T2, and T3 phenotypes, as in Figure 1B and C, are shown. Control, 1P and T2 cells are also shown in Figure 3B, and T1 cells in Figure S2B.

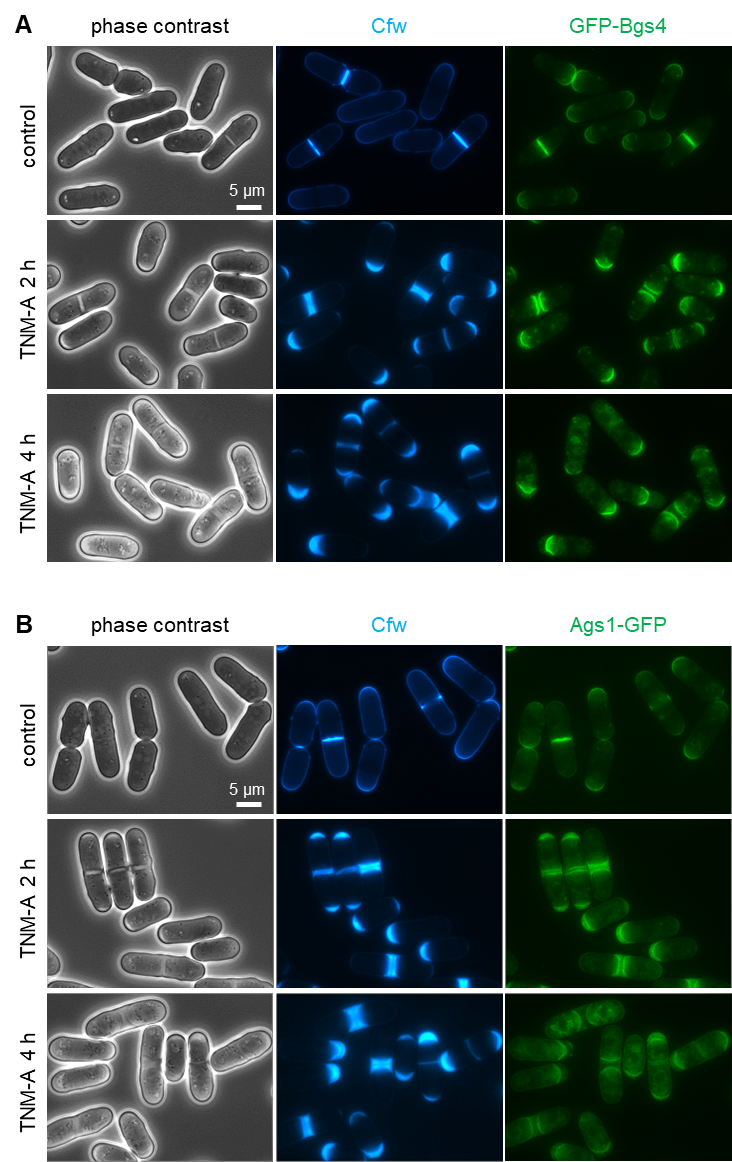

**Figure S8. Forced permanent localization of GFP-Bgs4 and Ags1-GFP by TNM-A.**

Localization of GFP-Bgs4 (A), or Ags1-GFP (B) in the presence of TNM-A. Cells expressing GFP-Bgs4, or Ags1-GFP were treated with TNM-A (2ug/mL). Images of phase contrast, and Cfw (mainly linear 1,3-β-glucan), and GS (GFP-Bgs4 in (A), Ags1-GFP in (B)) fluorescence are shown.

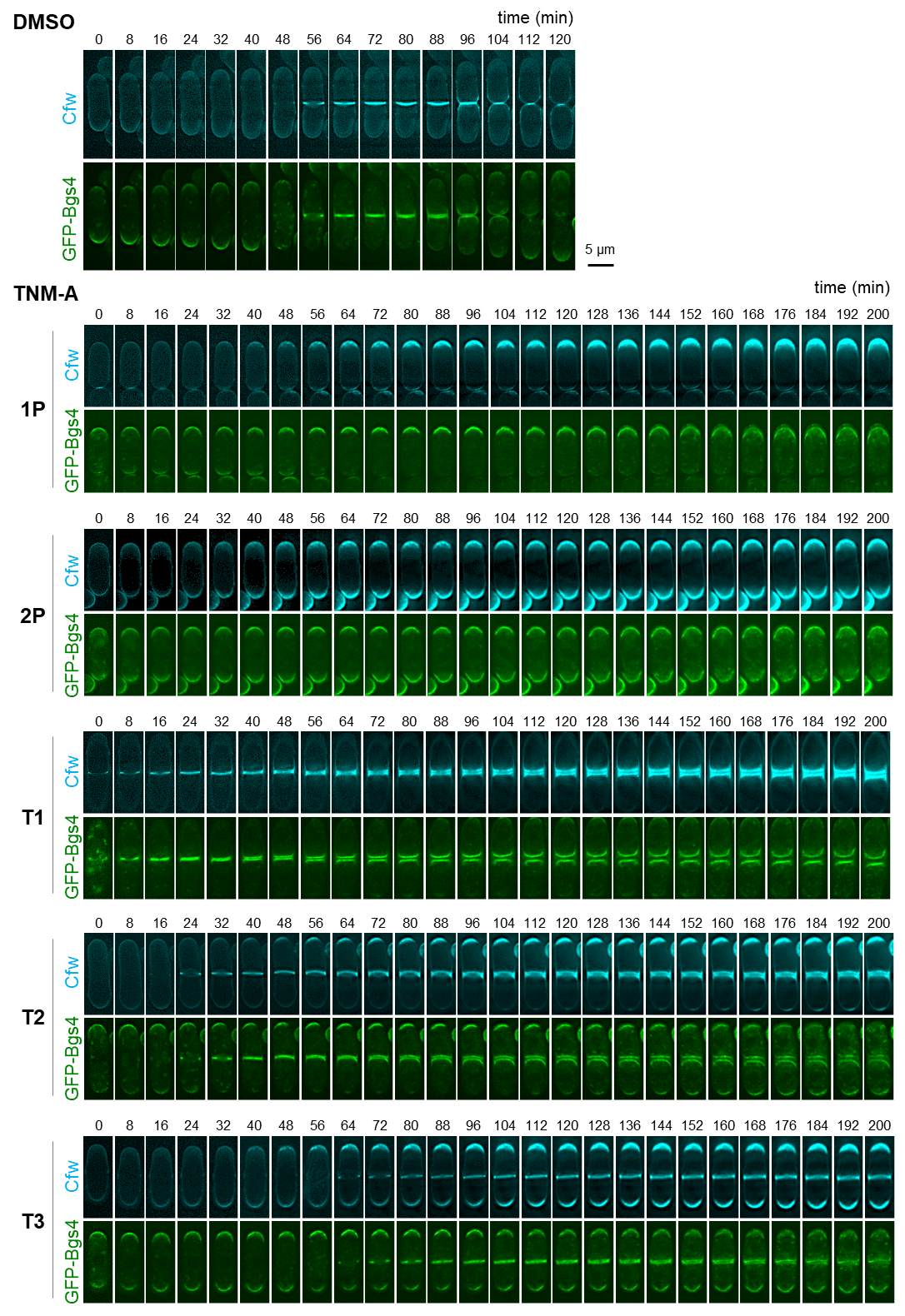

**Figure S9. Time lapse images of GFP-Bgs4 in the presence of TNM-A.**

Time lapse images of cells containing GFP-Bgs4. Cells were incubated in YES medium at 28 °C with DMSO (control) or TNM-A (2.0 µg/mL). Image capture was performed every 8 minutes for at least 120 minutes in different channels to observe Cfw staining (top, cyan) and GFP (bottom, green) fluorescence. Control cells and TNM-A treated cells with 1P, 2P, T1, T2, and T3 phenotypes, as in Figure 1B and C, are shown. Control, 1P and T2 cells are also shown in Figure 3B, and T1 cells in Figure S2B.

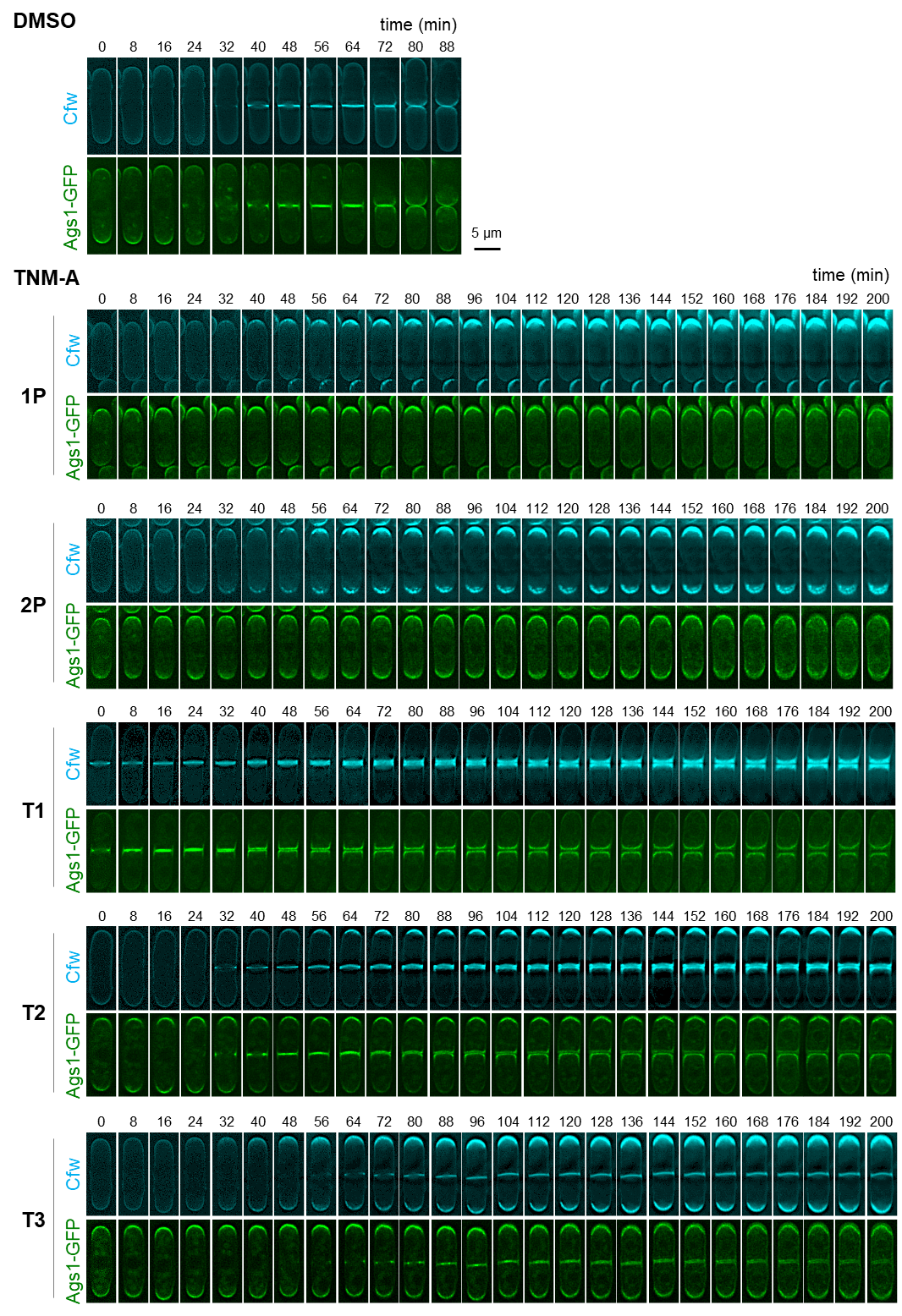

**Figure S10. Time lapse images of Ags1-GFP in the presence of TNM-A.**

Time lapse images of cells containing Ags1-GFP. Cells were incubated in YES medium at 28 °C with DMSO (control) or TNM-A (2.0 µg/mL). Image capture was performed every 8 minutes for at least 90 minutes in different channels to observe Cfw staining (top, cyan) and GFP (bottom, green) fluorescence. Control cells and TNM-A treated cells with 1P, 2P, T1, T2, and T3 phenotypes, as in Figure 1B and C, are shown. Control, 1P and T2 cells are also shown in Figure 3B, and T1 cells in Figure S2B.

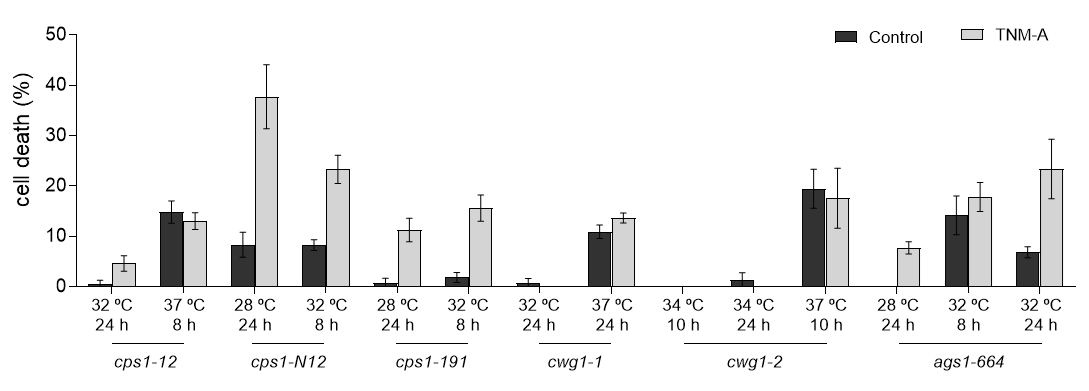

**Figure S11. Effect of TNM-A on the survival of GS mutant strains.**

GS mutant strains were grown in YES at 25 °C and were maintained in log phase. Then, each mutant was transferred to a semi-permissive and restrictive growth condition. The time indicates the maximum period each mutant can sustain at this temperature before growth stops. The last 4 hours of growth in semi-permissive or restrictive condition, TNM-A or DMSO (control) was added to the cultures. The wild-type strain is not shown, since it did not show cell death in any conditions. Data represent the mean ± SD (n = 3).

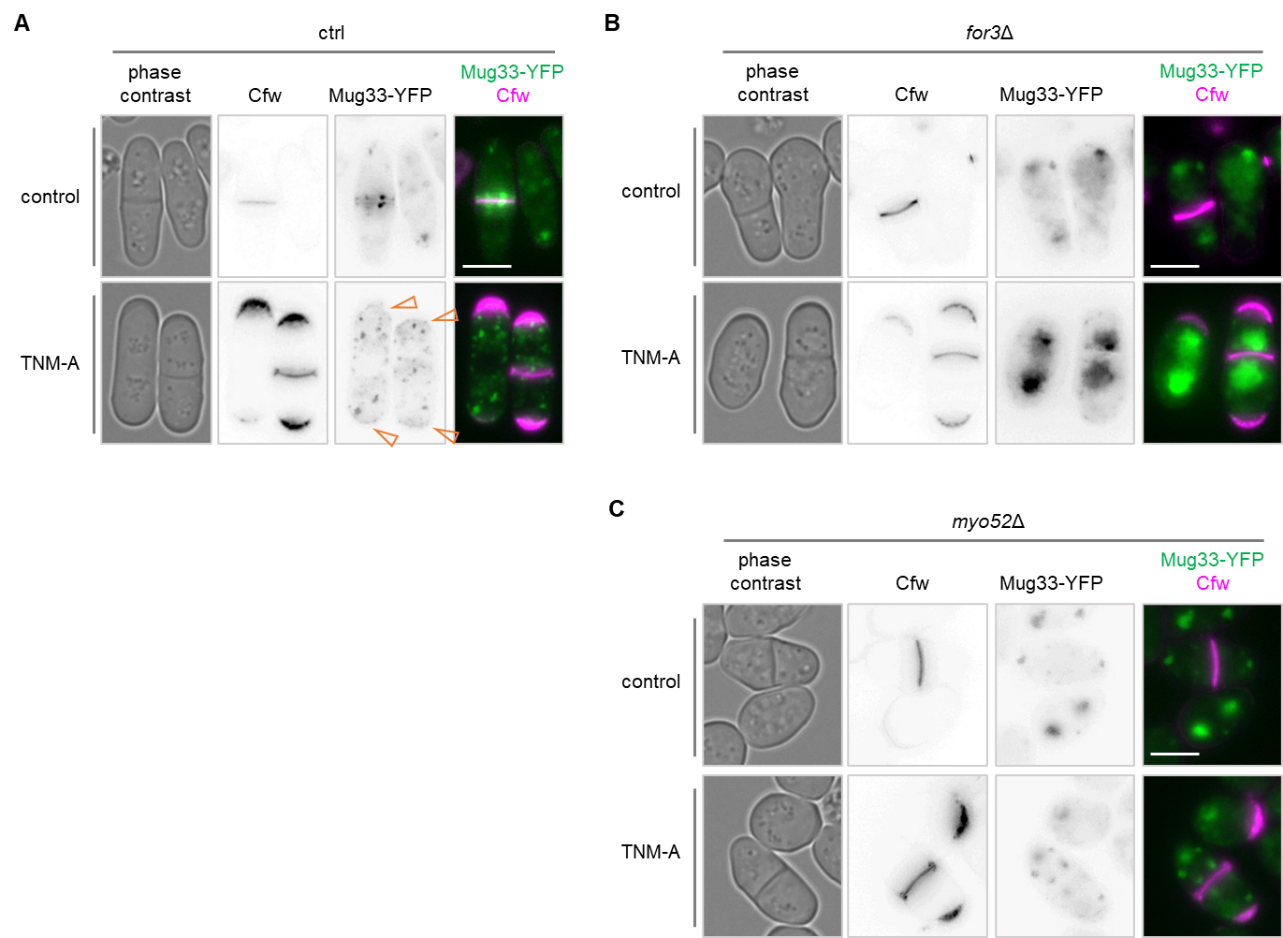

### **Figure S13. Effect of *for3*Δ or *myo52*Δ deletion on the localization of Mug33.**

Wild-type (A), *for3*Δ (B), or *myo52*Δ (C) cells expressing Mug33-YFP were treated with DMSO (control) or TNM-A (2 μg/ml) for 2 h. Cells were stained by Cfw (mainly linear 1,3-β-glucan) and observed under phase contrast and Cfw and YFP fluorescence microscopy. Open red triangle; accumulation of Mug33-YFP at the poles. Scale bar, 5 μm.

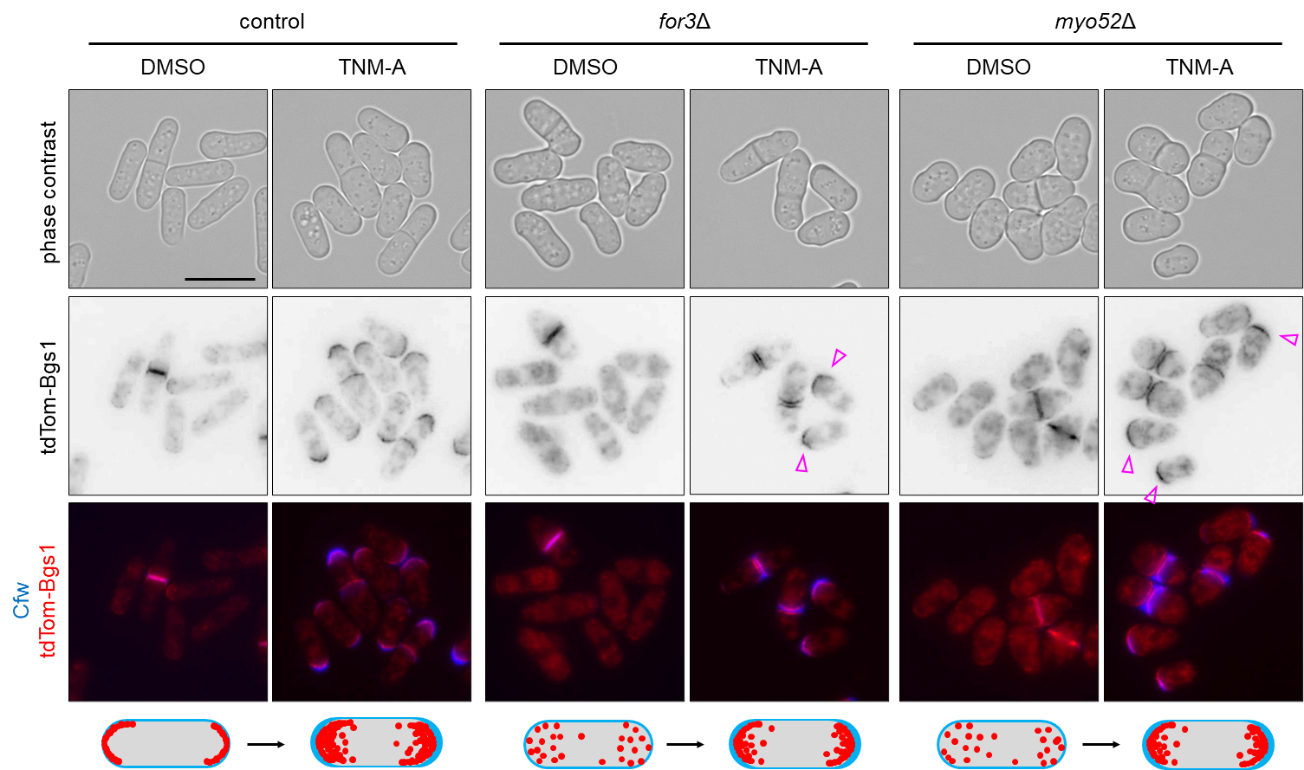

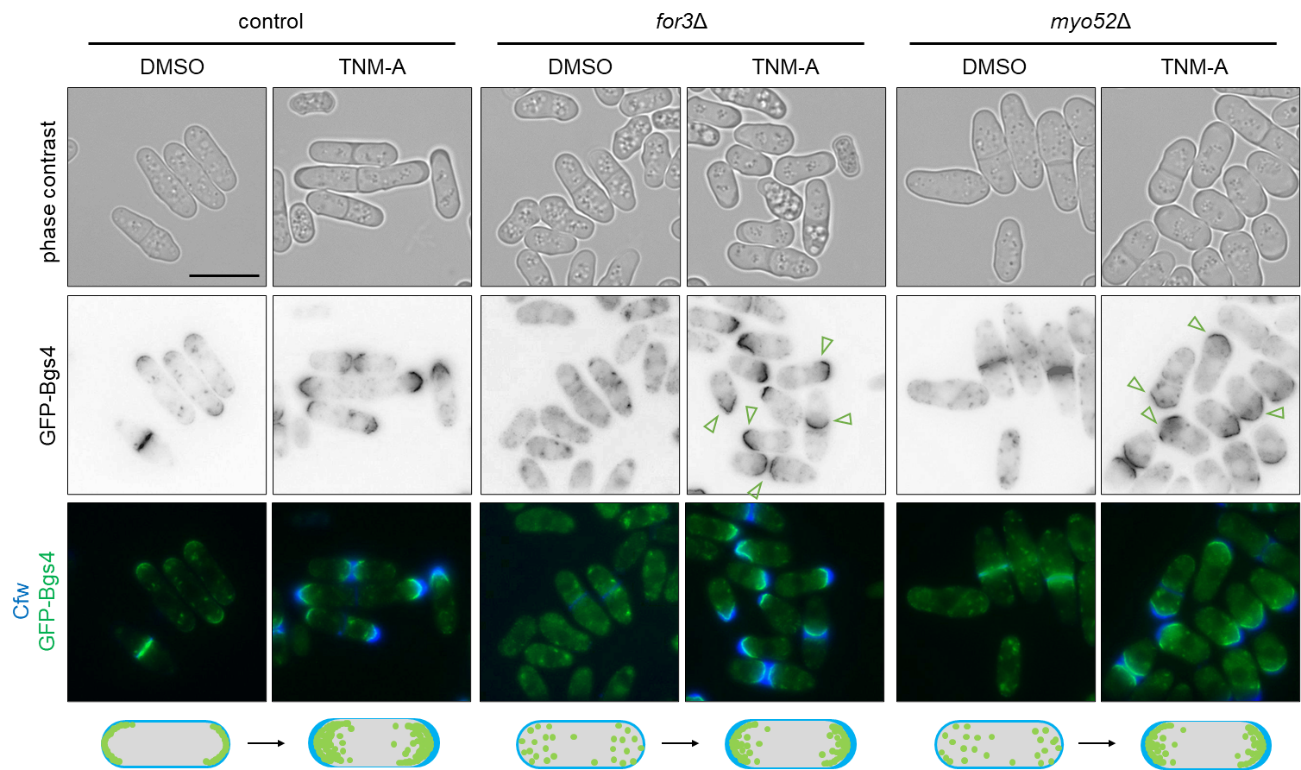

### **Figure S14. Effect of *for3* or *myo52* deletion on the localization of Bgs1, Bgs4, and Ags1.**

Control, *for3*Δ, or *myo52*Δ cells containing tdTom-Bgs1, GFP-Bgs4, or Ags1-GFP were treated with DMSO (control) or TNM-A (2 μg/ml) for 2 h. Cells were stained by Cfw (mainly linear 1,3-β-glucan) and observed under phase contrast and Cfw, tdTom, and GFP fluorescence microscopy. Scale bar, 10 μm. Open triangle; accumulation of tdTom-Bgs1 (red triangle), GFP-Bgs4, and Ags1-GFP (green triangle) at the poles. A scheme of the accumulation of Cfw, and tdTom-Bgs1, GFP-Bgs4, or Ags1-GFP at the poles with TNM-A is depicted bellow each series. Some of the pictures also appear in Figure 6.

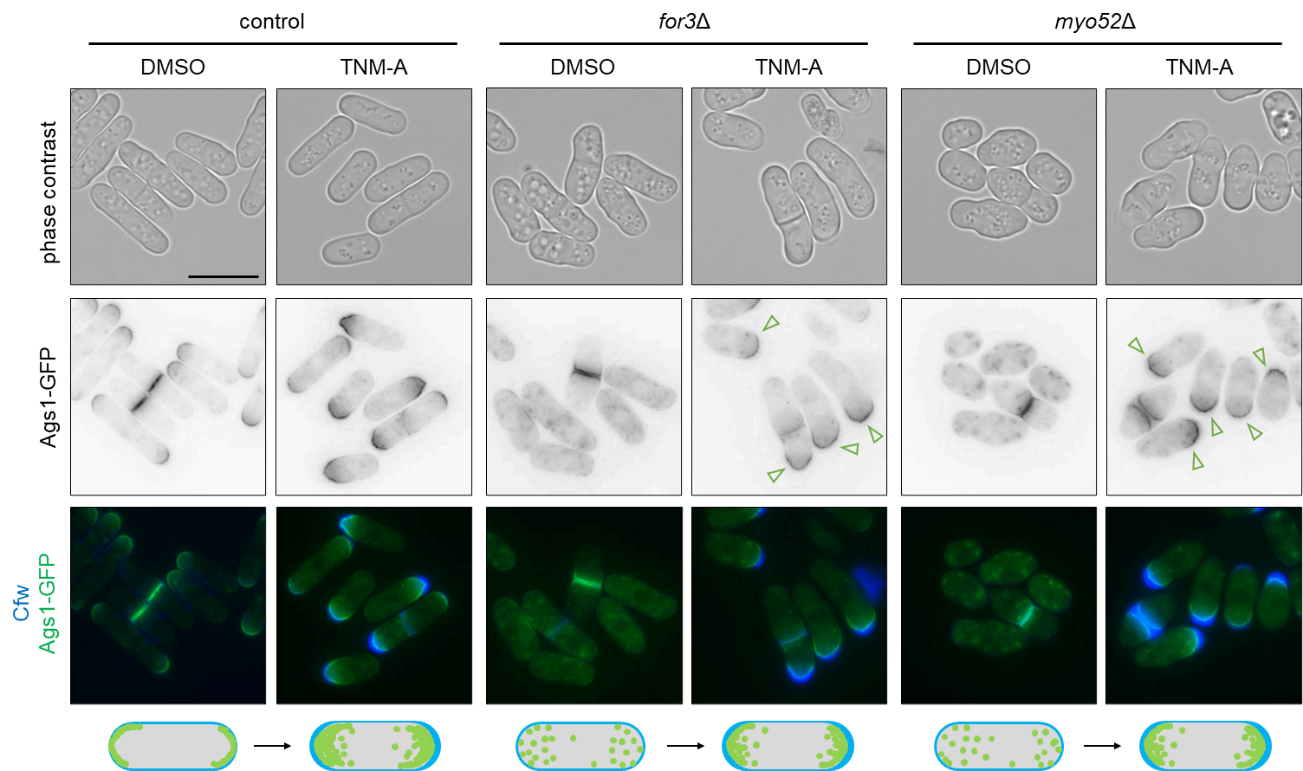

### **Figure S14. Effect of *for3* or *myo52* deletion on the localization of Bgs1, Bgs4, and Ags1 (continued).**

### **Table S1. List of 32 proteins that are localized at sites with thick cell wall in cells treated with TNM-A**

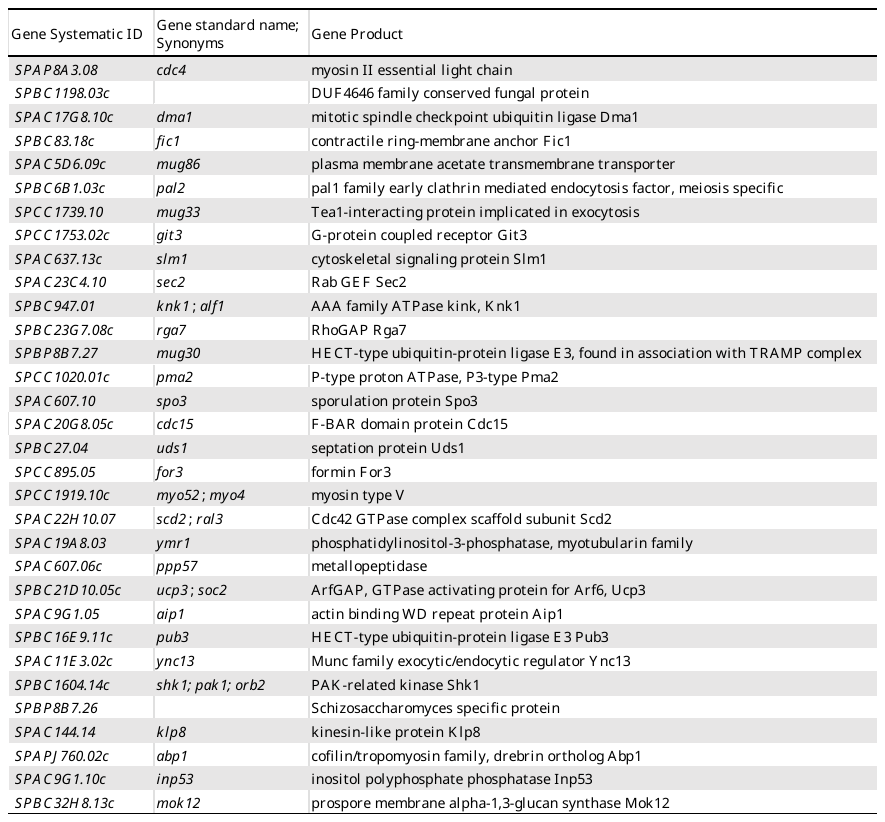
