## Supplementary Figure S12 for "Ectopic overproduction of cell wall glucan through membrane perturbation by an antifungal peptide theonellamide A in fission yeast"

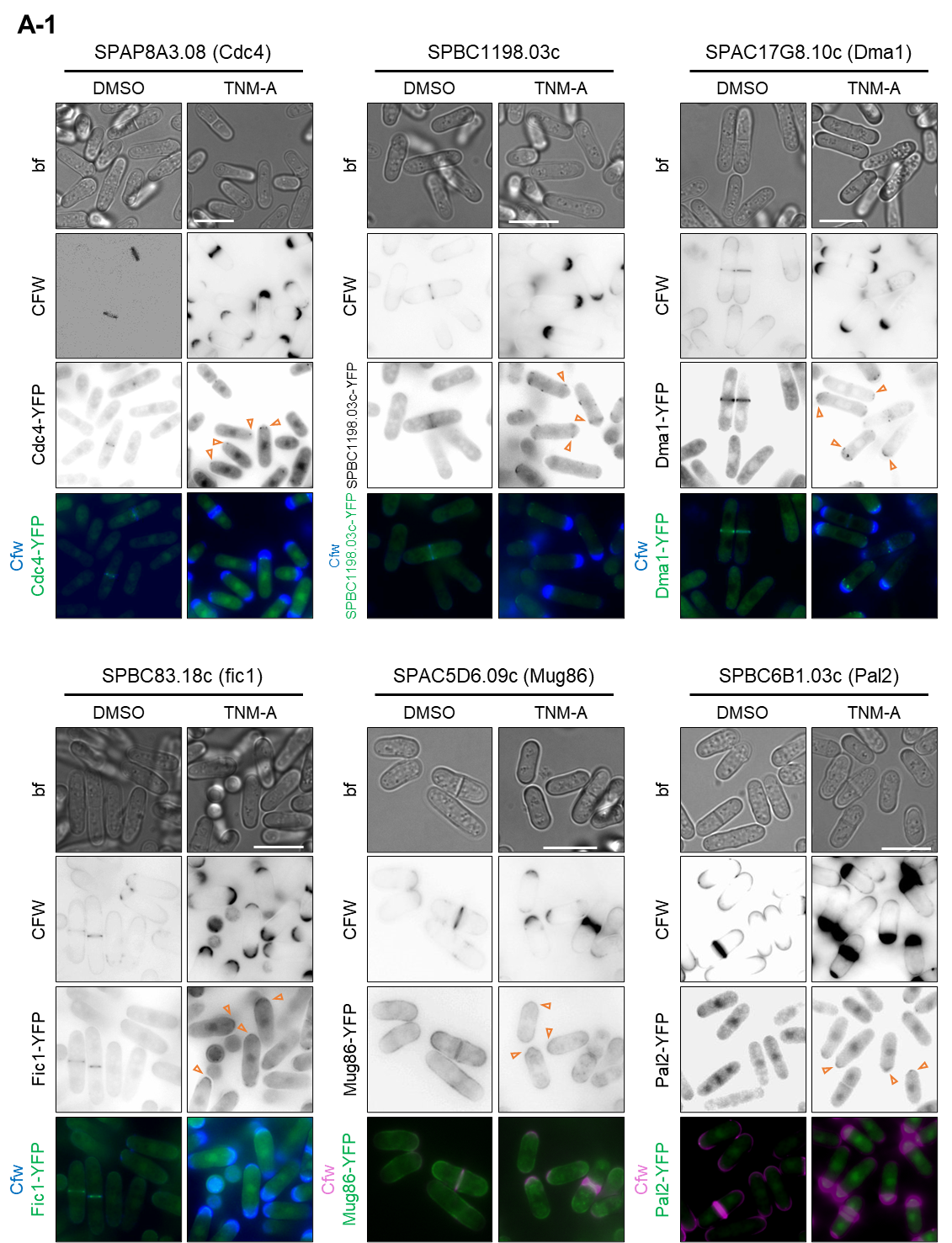

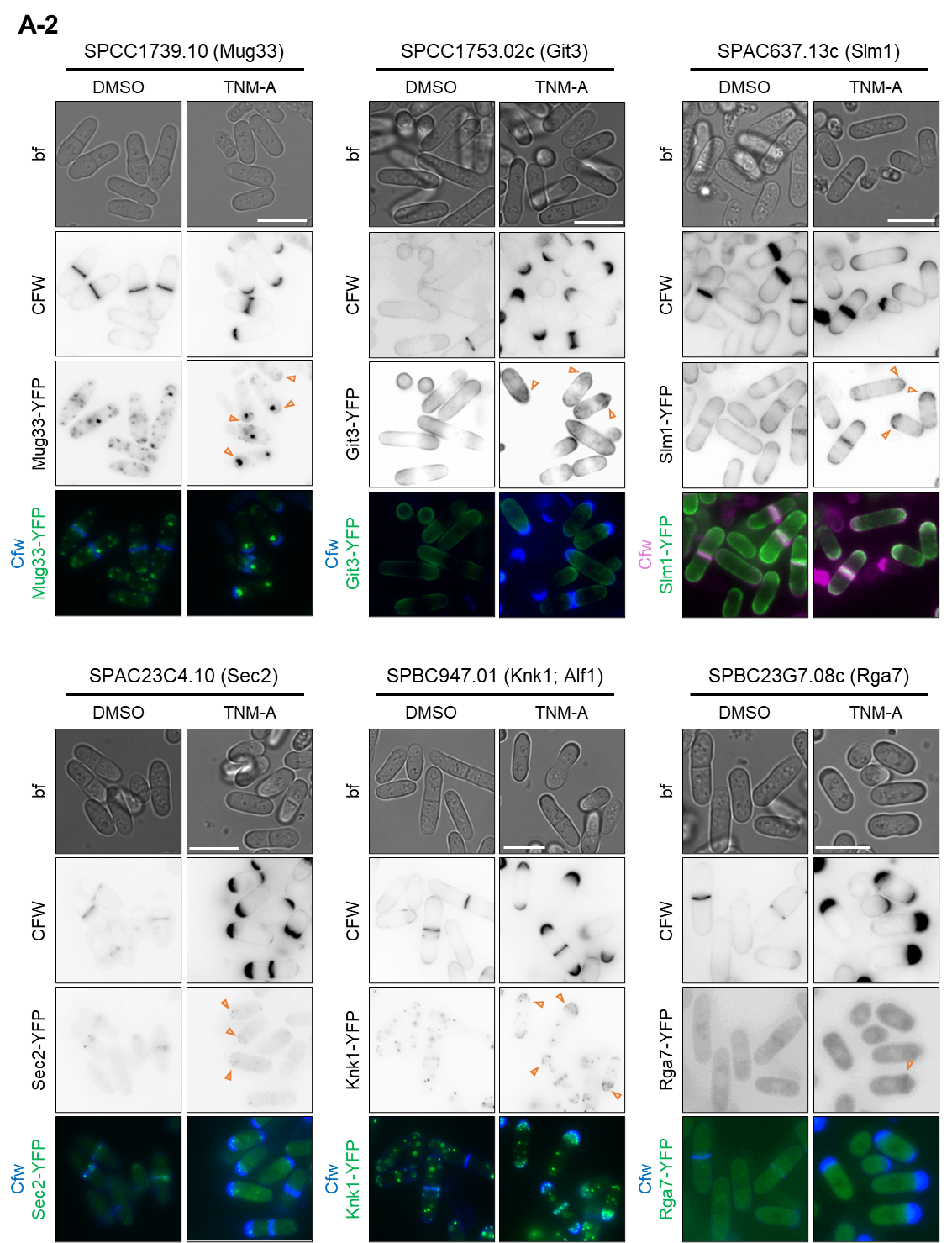

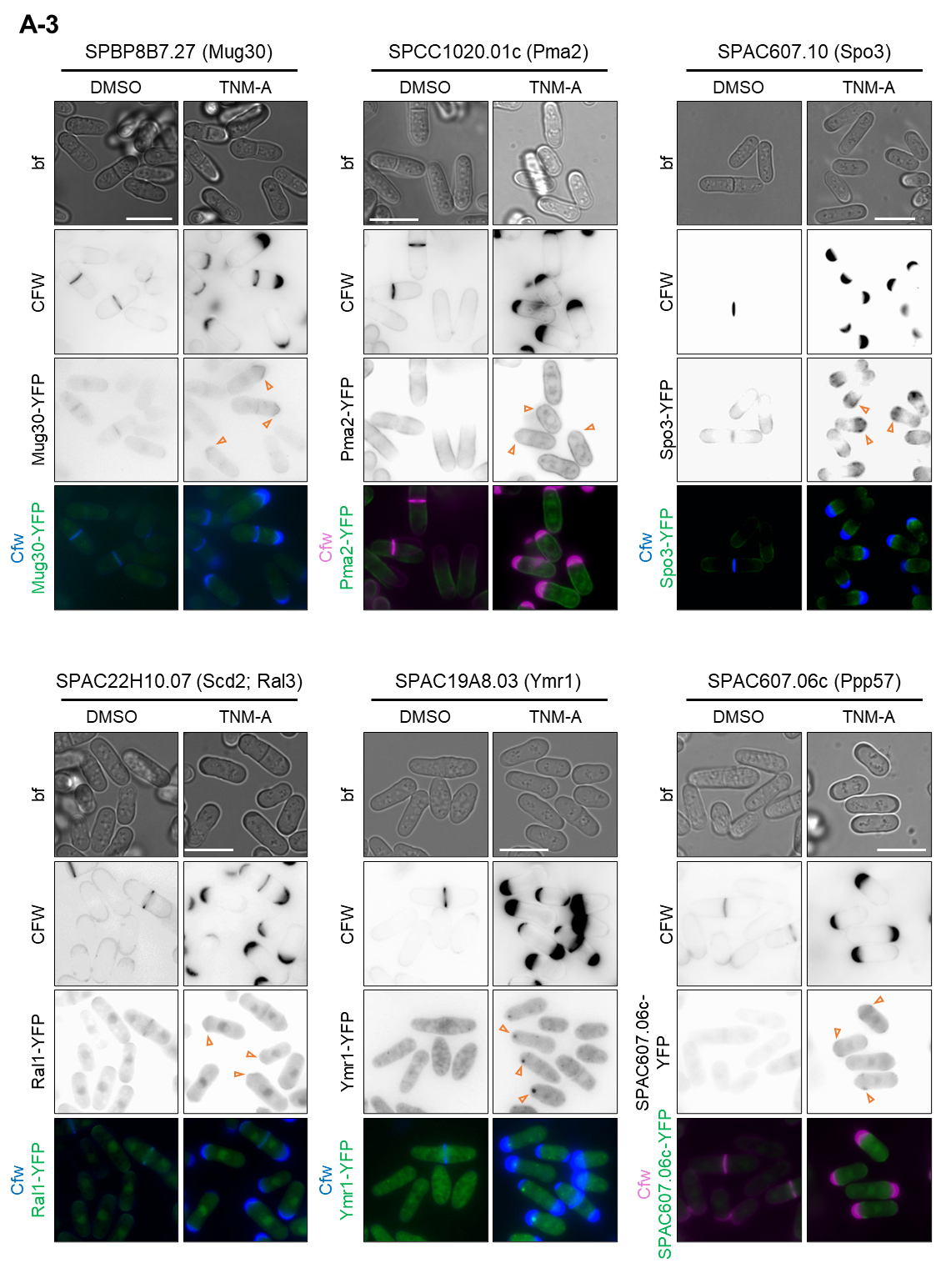

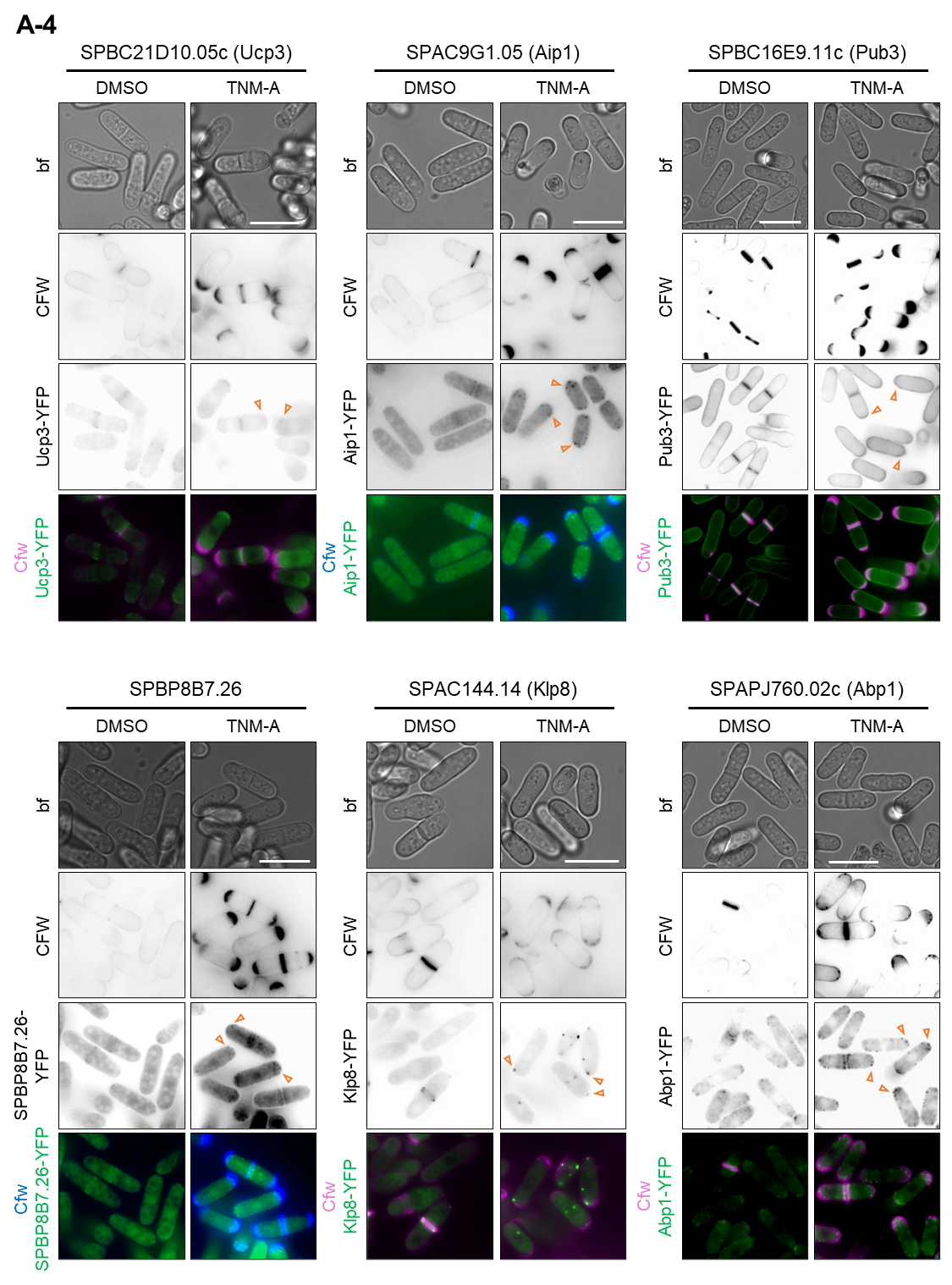

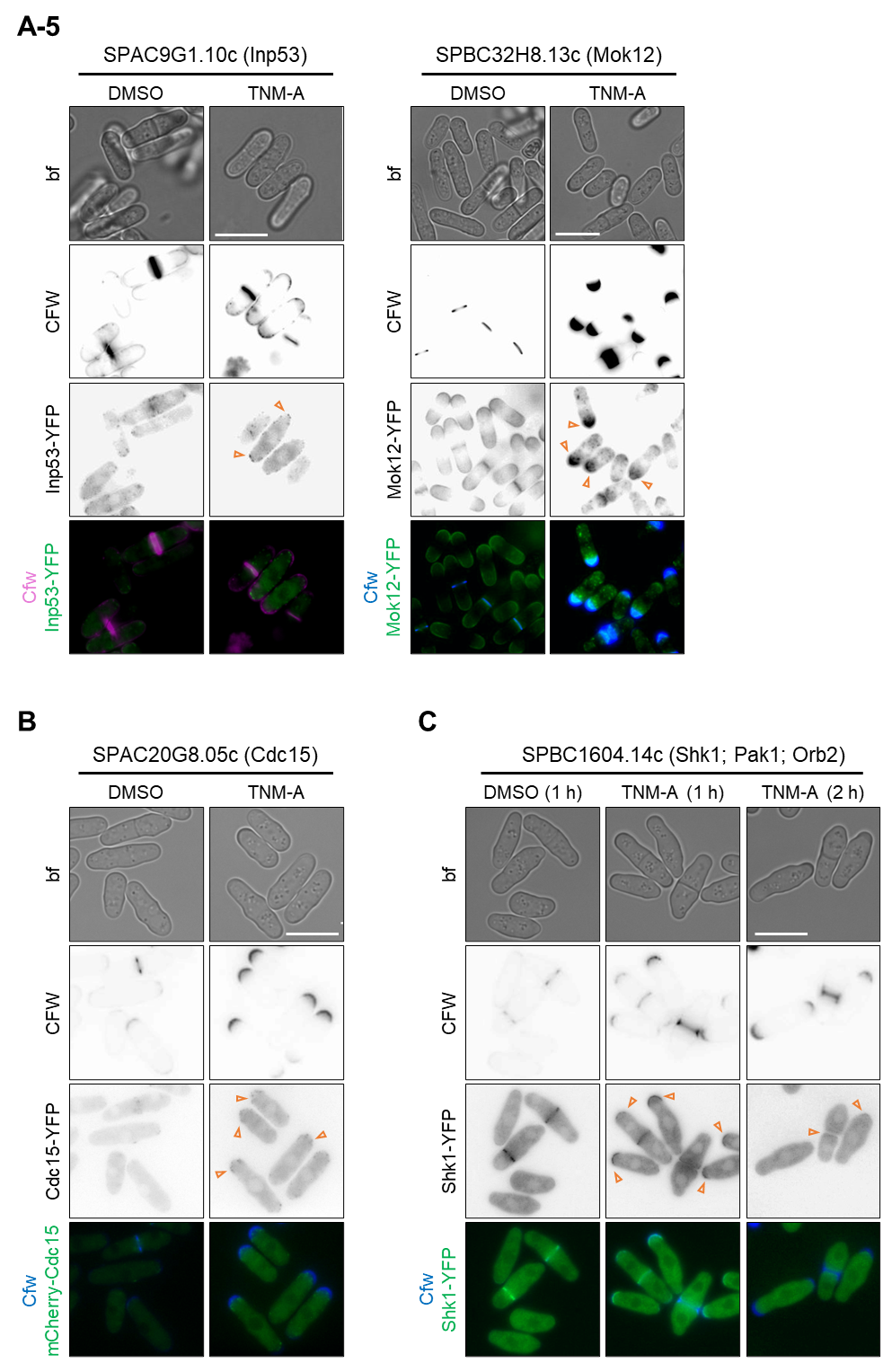

**Figure S12. Results of the localizome screen.** Subcellular localization of the proteins that are localized at/around thick cell wall are shown (orange arrowhead). Cells expressing YFP-tagged proteins under the control of the *nmt1* promoter or cells expressing mCherry-Cdc15 from the endogenous promoter were treated with DMSO or TNM-A (2 μg/ml) for 2 h (A1-A5, B). Cells expressing TFP-tagged Shk1 were challenged to DMSO or TNM-A (2 μg/ml) for 1 or 2 h. Cells were imaged for bright field (bf) and calcofluor white (CFW) and YFP fluorescence microscopy. Scale bars, 10 μm.
