## Supplementary Table S2 for "Ectopic overproduction of cell wall glucan through membrane perturbation by an antifungal peptide theonellamide A in fission yeast"

Table S2. *S. pombe* strains used in this work.

| strain | genotype | origin |
|---|---|---|
| JY1 | h$^{-}$ | lab collection |
| KGY16700 | h$^{+}$ mCherry-cdc15$^{+}$:kan$^{r}$ sid4$^{+}$-RFP:kanR ade6-M21X leu1-32 ura4-Δ18 | K.L. Gould [a] |
| KN2 | h$^{-}$ mCherry-cdc15$^{+}$:kan$^{r}$ | this study |
| CA5931 | h$^{-}$ leu1-32 ura4-294::[shk1 promoter:ScGIC2 CRIB-GFP3:ura4$^{+}$] | K. Shiozaki [b] |
| JCR3591 | h$^{+}$ leu1-32 ura4-Δ18 his3-Δ1 bgs1Δ::ura4$^{+}$ Pbgs1$^{+}$::tdTom-12A-bgs1$^{+}$:leu1$^{+}$ | lab collection |
| KN48 | h$^{-}$ sec3-913-HphR$^{ts}$ bgs1Δ::ura4$^{+}$ Pbgs1$^{+}$::tdTom-12A-bgs1$^{+}$:leu1$^{+}$ | this study |
| KN53 | h$^{-}$ sec3-913-HphR$^{ts}$ bgs4Δ::ura4$^{+}$ Pbgs4$^{+}$::GFP-12A-bgs4$^{+}$:leu1$^{+}$ | this study |
| KN13 | h$^{90}$ mug33$^{+}$-YFP for3Δ::kan$^{r}$ | this study |
| KN15 | h$^{90}$ mug33$^{+}$-YFP myo52Δ::kan$^{r}$ | this study |
| JCR3168 | h$^{-}$ leu1-32 ura4-Δ18 his3-Δ1 ags1Δ 3'UTRags1$^{+}$::ags1$^{+}$-12A-GFP-12A:leu1$^{+}$:ura4$^{+}$ | lab collection |
| KN18 | h$^{-}$ ags1Δ 3'UTRags1$^{+}$::ags1$^{+}$-12A-GFP-12A: leu1$^{+}$:ura4$^{+}$ | this study |
| KN21 | h$^{+}$ ags1Δ 3'UTRags1$^{+}$::ags1$^{+}$-12A-GFP-12A: leu1$^{+}$:ura4$^{+}$ for3Δ::kan$^{r}$ | this study |
| KN23 | h$^{-}$ ags1Δ 3'UTRags1$^{+}$::ags1$^{+}$-12A-GFP-12A: leu1$^{+}$:ura4$^{+}$ myo52Δ::kan$^{r}$ | this study |
| JCR1780 | h$^{-}$ leu1-32 ura4-Δ18 his3-Δ1 bgs1Δ::ura4$^{+}$ Pbgs1$^{+}$::tdTom-12A-bgs1$^{+}$:leu1$^{+}$ | lab collection |
| KN24 | h$^{-}$ bgs1Δ::ura4$^{+}$ Pbgs1$^{+}$::tdTom-12A-bgs1$^{+}$:leu1$^{+}$ | this study |
| KN26 | h$^{-}$ bgs1Δ::ura4$^{+}$ Pbgs1$^{+}$::tdTom-12A-bgs1$^{+}$:leu1$^{+}$ for3Δ::kan$^{r}$ | this study |
| KN28 | h$^{90}$ bgs1Δ::ura4$^{+}$ Pbgs1$^{+}$::tdTom-12A-bgs1$^{+}$:leu1$^{+}$ myo52Δ::kan$^{r}$ | this study |
| KN36 | h$^{-}$ bgs4Δ::ura4$^{+}$ Pbgs4$^{+}$::GFP-12A-bgs4$^{+}$:leu1$^{+}$ | this study |
| KN38 | h$^{+}$ bgs4Δ::ura4$^{+}$ Pbgs4$^{+}$::GFP-12A-bgs4$^{+}$:leu1$^{+}$ for3Δ::kan$^{r}$ | this study |
| KN41 | h$^{-}$ bgs4Δ::ura4$^{+}$ Pbgs4$^{+}$::GFP-12A-bgs4$^{+}$:leu1$^{+}$ myo52Δ::kan$^{r}$ | this study |
| JCR33 | h$^{-}$ 972 | P. Munz [c] |
| JCR77 | h$^{-}$ leu1-32 | lab collection |
| JCR284 | h$^{-}$ leu1-32 ura4-Δ18 his3-Δ1 | lab collection |
| JCR420 | h$^{+}$ leu1-32 ura4-Δ18 | lab collection |
| JCR1722 | h$^{-}$ leu1-32 ura4-Δ18 his3-Δ1 bgs1Δ::ura4$^{+}$ Pbgs1$^{+}$::GFP-12A-bgs1$^{+}$:leu1$^{+}$ | lab collection |
| JCR1723 | h$^{+}$ leu1-32 ura4-Δ18 his3-Δ1 bgs1Δ::ura4$^{+}$ Pbgs1$^{+}$::GFP-12A-bgs1$^{+}$:leu1$^{+}$ | lab collection |
| JCR2364 | h$^{-}$ leu1-32 ura4-Δ18 his3-Δ1 bgs4Δ::ura4$^{+}$ Pbgs4$^{+}$::GFP-12A-bgs4$^{+}$:leu1$^{+}$ | lab collection |
| JCR2365 | h$^{+}$ leu1-32 ura4-Δ18 his3-Δ1 bgs4Δ::ura4$^{+}$ Pbgs4$^{+}$::GFP-12A-bgs4$^{+}$:leu1$^{+}$ | lab collection |
| JCR3166 | h$^{-}$ leu1-32 ura4-Δ18 his3-Δ1 ade6-M210 ags1Δ 3'UTRags1$^{+}$::ags1$^{+}$-12A-GFP-12A:leu1$^{+}$:ura4$^{+}$ | lab collection |
| JCR3167 | h$^{+}$ leu1-32 ura4-Δ18 his3-Δ1 ade6-M210 ags1Δ 3'UTRags1$^{+}$::ags1$^{+}$-12A-GFP-12A:leu1$^{+}$:ura4$^{+}$ | lab collection |
| JCR4047 | h$^{-}$ leu1-32 ura4-Δ18 his3-Δ1 ade6-M210 hht1$^{+}$-RFP:KanMX6 | J. Cooper d |
| JCR4079 | h$^{-}$ leu1-32 ura4-Δ18 his3-Δ1 bgs1Δ::ura4$^{+}$ Pbgs1$^{+}$::GFP-12A-bgs1$^{+}$:leu1$^{+}$ hht1$^{+}$-RFP:KanMX6 | this study |
| JCR4080 | h$^{-}$ leu1-32 ura4-Δ18 his3-Δ1 bgs4Δ::ura4$^{+}$ Pbgs4$^{+}$::GFP-12A-bgs4$^{+}$:leu1$^{+}$ hht1$^{+}$-RFP:KanMX6 | this study |
| JCR4082 | h$^{+}$ leu1-32 ura4-Δ18 his3-Δ1 ade6-M210 ags1Δ 3'UTRags1$^{+}$::ags1$^{+}$-12A-GFP-12A:leu1$^{+}$:ura4$^{+}$ hht1$^{+}$-RFP:KanMX6 | lab collection |
| JCR5016 | h$^{90}$ leu1-32 GFP-psy1$^{+}$:leu1$^{+}$ | C. Shimoda e |
| JCR5062 | h$^{+}$ leu1-32 GFP-psy1$^{+}$:leu1$^{+}$ rlc1$^{+}$-tdTom:NatMX6 | this study |
| JCR5654 | h$^{-}$ leu1-32 ura4-Δ18 hht1$^{+}$-RFP:KanMX6 | this study |
| JCR6348 | h$^{+}$ leu1-32 ura4-Δ18 GFP-psy1$^{+}$:leu1$^{+}$ rlc1$^{+}$-tdTom:NatMX6 | this study |

a. Howard Hughes Medical Institute and Department of Cell and Developmental Biology, Vanderbilt University School of Medicine, Nashville, TN.
b. Division of Biological Science, Nara Institute of Science and Technology, Ikoma, Japan.
c. Institute of Cell Biology, University of Bern, Switzerland
d. Department of Biochemistry and Molecular Genetics, University of Colorado Anschutz Medical Campus, Aurora, USA.
e. Department of Biology, Graduate School of Science, Osaka City University, Osaka, Japan.
